# Community science designed ribosomes with beneficial phenotypes

**DOI:** 10.1101/2021.09.05.458952

**Authors:** Antje Krüger, Andrew M. Watkins, Roger Wellington-Oguri, Jonathan Romano, Camila Kofman, Alysse DeFoe, Yejun Kim, Jeff Anderson-Lee, Eli Fisker, Jill Townley, Eterna participants, Anne E. d’Aquino, Rhiju Das, Michael C. Jewett

## Abstract

Functional design of ribosomes with mutant ribosomal RNA (rRNA) could expand opportunities for understanding molecular translation, building cells from the bottom-up, and engineering ribosomes with altered capabilities. However, such efforts have been hampered by cell viability constraints, an enormous combinatorial sequence space, and limitations on large-scale, 3D design of RNA structures and functions. To address these challenges, we developed an integrated community science and experimental screening approach for rational design of ribosomes. This approach couples Eterna, an online video game that crowdsources RNA sequence design to community scientists in the form of puzzles, with *in vitro* ribosome synthesis, assembly, and translation in multiple design-build-test-learn cycles. We applied our framework to discover mutant rRNA sequences that improve protein synthesis *in vitro* and cell growth *in vivo*, relative to wild type ribosomes, under diverse environmental conditions. This work provides new insights into ribosome rRNA sequence-function relationships, with implications for synthetic biology.

## INTRODUCTION

The bacterial ribosome is composed of three distinct ribosomal RNAs (rRNAs) and more than 50 ribosomal proteins (rProteins) separated into small (30S) and large (50S) subunits^1–3^. It is responsible for the molecular translation of a defined genetic template into sequence-defined polymers of amino acids (i.e., proteins), with rRNA components facilitating messenger RNA (mRNA) decoding, accommodating amino acid substrates, catalyzing peptide bond formation, and excreting proteins from the exit tunnel. Motivated by the ribosome’s central role in controlling molecular translation, efforts are rapidly expanding to redesign, build, and repurpose ribosomes to facilitate understanding of ribosome assembly and function^2,4–7^, fill knowledge gaps in the origins of life^8–11^, and advance biotechnology^12–16^.

Efforts to modify or redesign the ribosome typically focus on creating rRNA mutants with assigned defects, enhanced functions, or altered capabilities. However, several bottlenecks have made altering the natural rRNA difficult. First, because the ribosome’s function is necessary for life, cell viability constrains the rRNA mutations that can be made as many mutations are dominantly lethal^17–22^. Cell-free approaches offer an alternative strategy, but cell-free built bacterial ribosomes, such as those from *Escherichia coli*, are not as active as wild type ribosomes^8,23,24^. Second, the mutational space is massive. For example, the 1,542-nucleotide long 16S and 2,904-nucleotide long 23S rRNAs of the *E. coli* ribosome are critical for function; thus, the theoretical sequence space for rRNA mutation (i.e., 4^4446^) is intractable to study experimentally. Third, the ribosome’s shape, physiochemical, and dynamic properties have been evolved over billions of years to build proteins with ~20 canonical α-amino acids, making redesign for a new function non-trivial^25^. Taken together, these features have resulted in a limited understanding of how to rationally design the structure and function of rRNA that makes up the ribosome.

Community science has emerged as a new approach to rationally design RNA structures and functions. This approach has advantages over typical computational RNA design methods, which are thwarted by the exponential complexity of minimizing the free energy of target RNA structures^26^. Eterna is an internet-scale community science game in which players ‘solve’ RNA secondary structure design puzzles subject to the constraints imposed by state-of-the-art thermodynamic energy models. In ‘lab challenges,’ players submit RNA puzzle ‘solutions’ and select a subset of these by voting, which then are synthesized and tested in research labs. Previously, Eterna has successfully targeted RNA design challenges inaccessible to other algorithms^27^. Despite recent progress in community science applied to RNAs and new methods for computational design of 3D RNA structure and function^28,29^, no RNA design efforts to date have approached the size or complexity of the entire bacterial ribosome.

In this work, we developed a design-build-test-learn (DBTL) approach, implemented through the Eterna platform, to create mutant ribosomes with improved rRNA secondary structure energetics. Specifically, our approach connects community scientists, university scientists, and game developers through three synergistic, interlocking, and mutually reinforcing DBTL cycles: game developers create 16S and 23S rRNA puzzles according to the needs of community scientists, community scientists design mutant 16S and 23S rRNAs by utilizing community and expert knowledge, and university scientists test these mutants by assessing them in an *in vitro* ribosome synthesis, assembly, and translation (iSAT) platform^8^. We demonstrate the power of our approach by conducting two iterations of a DBTL pipeline connecting three rounds of community science-derived ribosome design with a goal of improving protein expression in the iSAT cell-free platform. Through the course of this process, Eterna participants exceeded state-of-the-art computational predictions and designed mutant ribosomes with 42 ± 10% greater protein expression than wild type ribosomes under optimal conditions and across diverse stress conditions *in vitro*. Surprisingly, the mutant ribosomes also support life and enable faster growth than cells grown exclusively with wild type ribosomes. We anticipate that our Eterna-based DBTL approach will be valuable for engineering complex RNA machines, advancing our knowledge about RNA sequence-folding-function relationships, and inspiring new directions to engineer ribosomes for synthetic biology applications.

## RESULTS

In this study, we aimed to demonstrate the use of the Eterna platform to crowdsource the design of ribosomes with functional, stabilized mutant rRNAs with beneficial phenotypes. Following a “pilot round” (R0), where we discovered Eterna players could design functional ribosomes with unexpectedly large numbers of rRNA mutations, we devised a DBTL framework for iterative citizen science collaborations and applied it to two further rounds of the “OpenRibosome Challenge.”

### Eterna participants can design mutant ribosomes with diverse sequences that outperform computationally predicted designs

We aimed to explore the use of the Eterna platform to crowdsource the design of functional 16S and 23S rRNA variants with improved secondary structure energetics, as compared to the wild type (WT) *E. coli* sequences. For this, we first carried out a “pilot round” (R0) of puzzles on the Eterna platform for the ribosome’s 16S and 23S rRNAs and used a conventional Design-Build-Test-Learn (DBTL) strategy (Fig. 1a). In the <u>D</u>esign phase, the Eterna platform released 16S and 23S rRNA puzzles, and participants designed mutant rRNA sequences by exchanging individual nucleotides against any other unmodified RNA nucleotide (A, C, G, U) in the puzzles, with some critical nucleotides “locked” to their WT identities (see Methods). These sequences then were scored with a folding engine calculating the free energy (ΔG) of the sequence’s secondary structure (see Methods, “Design of ribosome puzzles”) and submitted to a pool of designs from which the players then voted on the best 8 designs. In the Build phase, we synthesized plasmid DNA encoding the rRNA designs selected by the players. In the <u>T</u>est phase, mutant 16S and 23S rRNAs were assessed in an *in vitro* ribosome construction platform, called iSAT^8,24,30^. iSAT enables one-pot co-activation of rRNA transcription, assembly of rRNA with native ribosomal proteins into *E. coli* ribosomes, and synthesis of functional proteins from these ribosomes in a crude S150 extract lacking native ribosomes (**Figure 1a**, bottom). A key feature of this system is the ability to generate ribosomal variants by simply changing the DNA input, which enables rapid screening of *E. coli* ribosome mutations^31,32^. Moreover, because ribosomes assembled in iSAT have lower activity than *in vivo*-assembled versions^8,23,24^ and there are known inefficiencies with ribosome reconstitution *in vitro*^8,23^, we hypothesized that iSAT would enable us to identify ribosomes where stabilized rRNAs could lead to improved activity. Finally, in the <u>L</u>earn phase, we shared the results with the players on the Eterna webpage in the form of detailed results postings.

**Figure 1:**
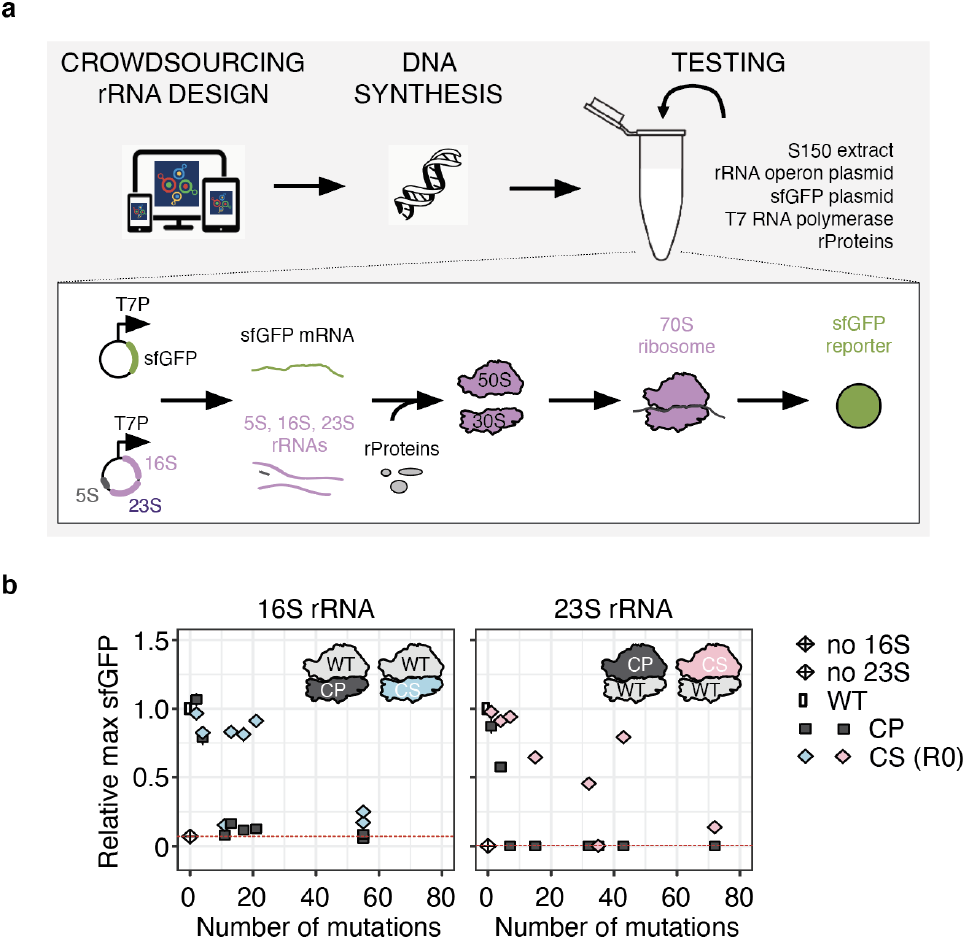
Crowdsourcing rRNA design enables functional mutant ribosomes and outperforms computational predictions. (a) rRNA design was crowdsourced to community scientists (CS) via the Eterna platform. In a “pilot round” (R0), CS solved 16S rRNA and 23S rRNA puzzles, submitted their “solutions” and voted on 8 designs per puzzle. These designs were synthesized and tested for activity *in vitro* by integrated ribosome synthesis, assembly, and translation (iSAT). (b) Comparison of sfGFP expression in iSAT reactions with pT7-rrnB-16S and pT7-rrnB-23S rRNA variants generated either by computationally prediction (CP or by CS). sfGFP expression was determined by fluorescence over 16 hours and normalized to the maximum sfGFP of pT7-rrnB-wild type (WT). Data are shown as mean ± s.d.; n ≥ 3. Dotted red line indicates background activity of the S150 extract due to residual ribosomal subunits.

Seventeen players submitted 129 16S rRNA designs and sixteen players submitted 157 23S rRNA designs. The players then voted on their top eight designs for each rRNA. The 16S rRNA designs comprised 2-55 mutations and 23S rRNA designs harbored 1-72 mutations distributed over the entire rRNA sequences (**Suppl. Data 1; Suppl. Table 1**). DNA sequences were then synthesized and cloned in place of the corresponding 16S rRNA or 23S rRNA into plasmid pT7-rrnB, which encodes a copy of the *rrnB* operon controlled by the T7 promoter, tested in the iSAT platform, and results shared with the Eterna community (**Figure 1a**). Six of the 16S rRNA and seven of the 23S rRNA designs from the players were functional in iSAT, conferring activities between 14 and 96% compared to WT with only a mild negative correlation between number of 16S/23S rRNA mutations and iSAT activity. We compared the iSAT activity of the 16S and 23S rRNA variants designed by community scientists (CS) head-to-head with a set of computationally predicted (CP) designs (see Methods) with exactly matched mutation numbers (**Figure 1b; Suppl. Figure 1**) and subject to the same secondary structure requirements. We found that CS designs were able to maintain nearly WT-like performance despite installing >20 mutations (16S rRNA) or even >40 mutations (23S rRNA), while every CP design with more than 4 mutations was inactive in iSAT. While we have previously used iSAT to show that more than 85% of 180 single nucleotide mutations within the ribosome active site possess some functional activity^31^, we were surprised that so many mutations could be designed into active ribosomes.

To determine the robustness of the initial Eterna 16S rRNA and 23S rRNA designs, we took advantage of the iSAT platform’s direct access to the reaction environment to perturb the magnesium (Mg^2+^) concentration, which impacts rRNA folding and intrinsic rRNA folding stability^33,34^. For this, we set up iSAT reactions with half of the experimentally optimized Mg^2+^ concentration (3.75 mM instead of 7.5 mM) and compared the activity of CS-designed rRNA variants with WT rRNA. While WT ribosome activity was still highest with 42% activity compared to optimal iSAT conditions, most CS-designed rRNAs assembled into functional ribosomes at low Mg^2+^ concentration, leading to activities of up to 41% of WT (**Suppl. Figure 2**). These results show that the CS rRNA designs, which were implemented on the basis of RNA secondary structure folding energetics, are robust under non-optimal iSAT conditions. In sum, the “pilot round” allowed us to build a community of Eterna participants with interest in ribosome design challenges, implement puzzles for rRNAs in the Eterna design framework, and realize that community scientists can outperform computational prediction methods in designing functional 16S and 23S rRNAs by balancing numerous constraints in the design process and selecting the best designs through the voting process.

### Development of a progressive design-build-test-learn pipeline for designing diverse 16S and 23S rRNAs

Following the pilot round, we recognized that continuing our initial DBTL approach would limit the project’s progress. While the size and complexity of the rRNA molecules and the assay used to test them excited Eterna players, they were also overwhelming. The puzzles were difficult to play through the Eterna interface because the algorithm to display their structures would place multiple nucleotides at the same point in 2D space, making it impossible to mutate certain bases. Finally, the feedback available to players was limited to energetics calculated within the puzzles and the experimental results available months later, limiting their ability to explore diverse solution strategies. To this end, we realized that the Eterna platform itself—from the ribosome puzzles to the analysis tools offered by the game—would require DBTL iteration in parallel, and that the same three parties—university scientists, game developers, and community scientists—would have to collaborate to advance an approach in which each of these typically disparate groups would work together to propel the science and gameplay in tandem (**Figure 2a**).

**Figure 2:**
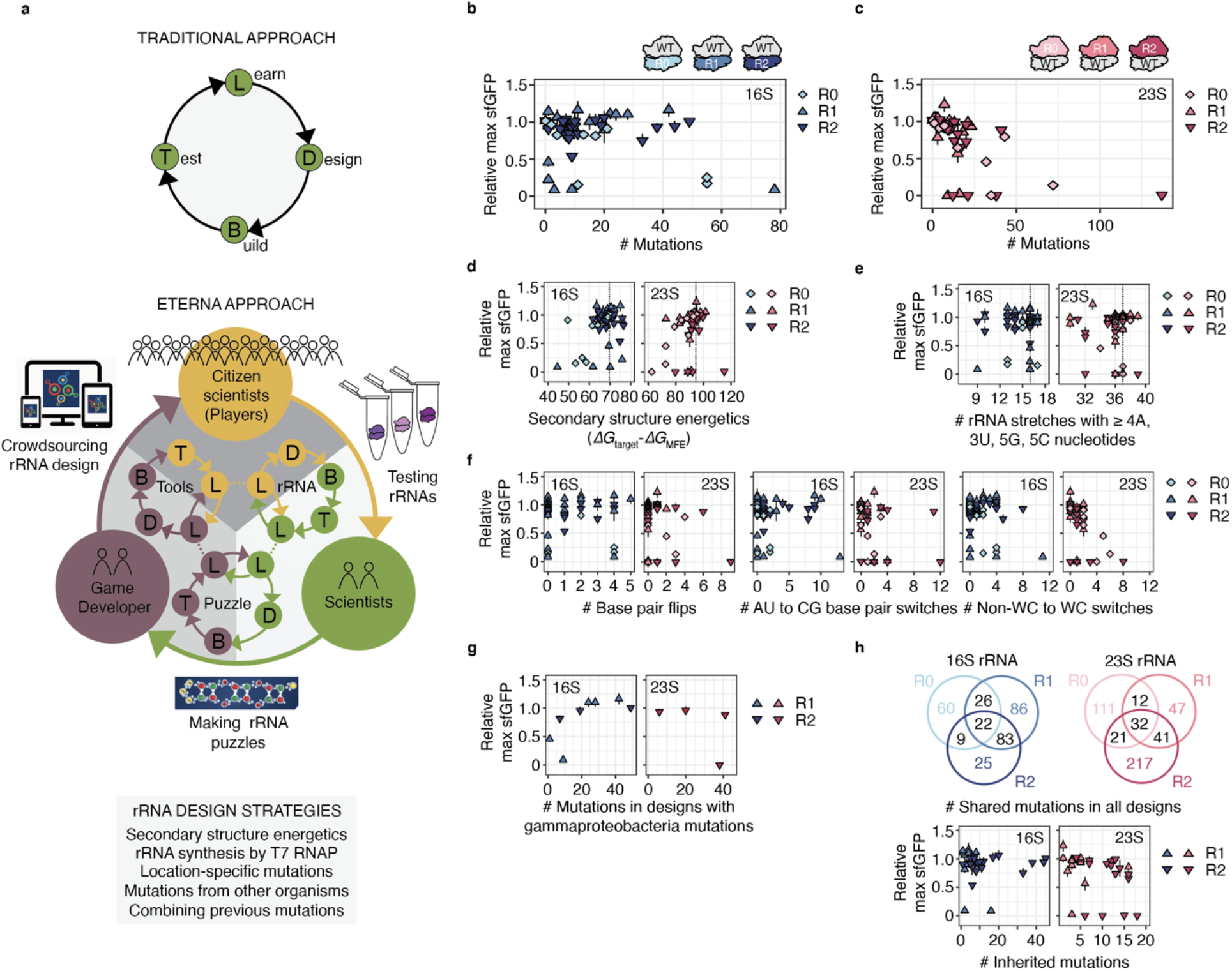
The Eterna approach transforms traditional design-build-test-learn cycles. (a) Our Eterna approach exploits community scientists (CSs) who solve RNA design challenges by closely working together with each other, university scientists, and game developers. Each of these parties contribute to synergistic, interlocking, and mutually reinforcing DBTL cycles, thereby boosting the iteration of making rRNA puzzles, crowdsourcing rRNA design to community scientists through the Eterna platform, and discovering rRNA variants in research labs. We performed two rounds of our approach. Functionality comparison of Round 1 (R1) and Round 2 (R2) Eterna 16S rRNA (b) and 23S rRNA (c) designs in iSAT with the pilot round (R0) in dependence of the designs’ mutation counts. In order to approach the challenge, CSs followed and combined different strategies: (d) secondary structure energetics, (e) breaking stretches of consecutive identical nucleotides, (f) altering base pairing in rRNA secondary structures, (g) integrating mutations from other gammaproteobacteria, and (h) integrating/ combining of mutations from previous rounds. sfGFP expression in iSAT was determined by fluorescence and normalized to max sfGFP of pT7-rrnB-wild type. Dotted line in (d) and (e) indicates the WT value. Data are shown as mean ± s.d.; n ≥ 3.

To test this framework, we set up a multi-round “OpenRibosome Challenge” on the Eterna platform with the aim of generating further stabilized rRNAs that lead to improved iSAT activity. We improved upon R0 in two notable ways. First, we programmed new software for RNA layout (https://github.com/ribokit/RiboDraw) and composed a diagram of the 16S and 23S rRNAs that reflects the relative position of elements in 3D space to avoid the overlapping rRNA sequences observed in the R0 puzzles. Second, we released two constraints over “locked” nucleotides. We made this change because during R0 we found that the “locked” nucleotides constrained player creativity, and after examining sequence variation across all gammaproteobacteria we noted that some residues deemed immutable were not conserved. The Eterna players developed an in-game system for tracking which mutations violated gammaproteobacteria sequence conservation (see Methods, “Design of ribosome puzzles”); we anticipated that this system would enable fine-grained feedback, leading to greater freedom to make extensive but high-confidence mutations than was possible with “locked” nucleotides.

We conducted two rounds of the OpenRibosome Challenge. In Round 1 (R1), a total of 31 players submitted 205 16S rRNA designs and 22 players submitted 161 23S rRNA designs. In Round 2 (R2), a total of 23 players submitted 139 16S rRNA designs and 20 players submitted 113 23S rRNA designs. During each round, the players selected 20 designs each of 16S rRNA and 23S rRNA, which then were synthesized and tested at optimal iSAT conditions (**Suppl. Data 1; Suppl. Table 1).** After each round, the data were shared with the community in the form of posts and discussed in forums and community meetings. We found that Eterna players learned from the iSAT data provided and improved their designs over time. For example, player designs in R1 and R2 had higher activity and more mutations than the R0 “pilot” 16S and 23S rRNA designs (**Figure 2b–c; Suppl. Figure 3**).

Several sequence-function strategies emerged for designing stabilized rRNAs. These included: optimizing secondary structure energetics to support intrinsic rRNA folding, especially in R0 (**Figure 2d**), and minimizing sequence repeats to prevent T7 RNA polymerase-derived rRNA synthesis errors (**Figure 2e; Suppl. Figure 4a**). When making mutations, players predominantly flipped base pairs, switched AU to CG pairs and vice versa, and wobble (non-Watson-Crick (non-WC)) base pairs to WC base pairs (**Figure 2f; Suppl. Figure 4b**). Furthermore, guided by scientific literature, players also successfully incorporated mutations from other gammaproteobacteria, with a focus on extremophilic bacteria that are known to have favorable RNA folding capabilities^4,5^ (**Figure 2g**). The players used designs from current or previous rounds as inspiration, borrowing and recombining mutations with great success (**Figure 2h**). In addition, players found it beneficial to prevent exchanging nucleotides that are conserved or directly contact rProteins^35,36^ (**Suppl. Figure 4c**) and they avoided previously described deleterious mutations^21,22,31^.

To characterize and identify robust Eterna rRNA designs, we selected all R1 and R2 designs with ≥ 6 mutations that showed activity of ≥ 80% WT at optimal iSAT conditions as a curated set of high performing, high sequence diversity ribosomes and tested them under folding stress conditions: low <u>M</u>g^2+^ (3.75 mM) concentration and <u>o</u>ptimal temperature (37 °C), LMOT; or low <u>M</u>g^2+^ (3.75 mM) concentration and low temperature (30 °C), LMLT (**Figure 3a; Suppl. Figure 5-6; Suppl. Table 1**). Several Eterna designs showed robustness to these folding stress conditions, especially the R1 16S designs.

**Figure 3:**
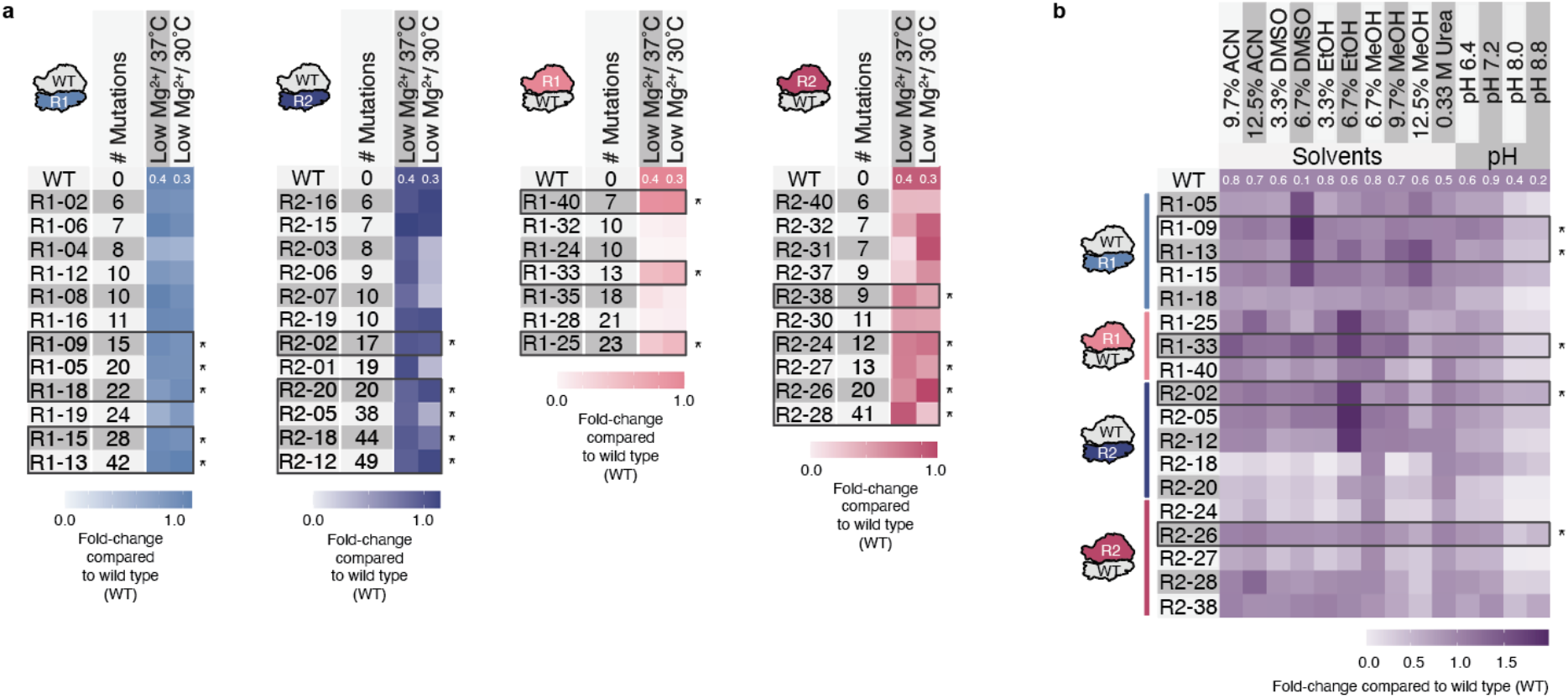
Eterna rRNA designs are robust across diverse stress conditions *in vitro*. Designs with ≥ 6 mutations and an activity of ≥ 80% wild type (WT) at optimal iSAT conditions were tested in diverse iSAT stress conditions. (a) Heatmaps illustrating folding stress tolerance of Eterna R1 and R2 16S and 23S designs with ≥ 6 mutations and an activity at optimal iSAT conditions of ≥ 80 % WT in iSAT. (b) Heatmap illustrating solvent and pH tolerance of designs with robust folding phenotypes and high mutation rate in iSAT. sfGFP expression in iSAT was determined by fluorescence and normalized to the maximum sfGFP of pT7-rrnB-wild type at optimal iSAT conditions. Data are shown as mean; n ≥ 3. ACN: acetonitrile, DMSO: dimethylsulfoxide, EtOH: ethanol, MeOH: methanol. As reference, activities of WT at each iSAT stress condition compared to iSAT at optimal conditions and the number of mutations of the most mutated design are provided as white numbers in each heatmap. * selected in (a) as designs for being tested in (b), in (b) as designs with outstanding features (i.e., high iSAT activity in multiple stress conditions).

From this set of 41 designs, we identified the 18 most diverse (i.e., high mutation rate) and robust (i.e., high iSAT activity) designs per rRNA and round. These 18 designs were next tested in ISAT under non-physiological conditions; the presence of organic solvents (ACN: acetonitrile, DMSO: dimethylsulfoxide, MeOH: methanol, EtOH: ethanol) or altered pH (**Figure 3b; Suppl. Figure 8-12**). We selected solvent conditions that reduce iSAT activity using WT rRNAs (**Suppl. Figure 7**). Strikingly, several Eterna-designed ribosomes from R1 and R2 exceeded WT ribosome performance, including R1-13 and R1-15 at 9.7% MeOH, R1-09 at 12.5% MeOH, R2-02 at 6.7% EtOH, as well as R1-33 and R2-26 at 12.5% ACN (**Figure 3b; Suppl. Figure 8-12**). Together, these data highlight the potential for community science to develop ribosomes with beneficial phenotypes, and how computational design might be used to build modified ribosomes with altered chemical properties in the future.

### Combining Eterna 16S and 23S rRNA designs increases in vitro ribosome activity

We next combined orthogonal Eterna player designs to assess the impact on *in vitro* ribosome synthesis and activity. We selected three 16S rRNA designs (R1-09, R1-13, R2-02) and two 23S rRNA (R1-33, R2-26) designs covering both rounds. These designs were selected because they had the best combination of sequence diversity, optimal iSAT performance, and tolerance to iSAT stress conditions. After building plasmids encoding the combined designs, we tested the six combinations (each designed small subunit paired with a designed large subunit) in iSAT reactions (**Figure 4; Suppl. Figure 13, 14**). In iSAT at optimal conditions, the R1-09/R1-33 combination, totaling 28 mutations, showed 142 ± 10 % activity, a significantly higher level of function than the WT (**Figure 4a**). This advantage was also observed in several solvent stress conditions (**Figure 4b**). Under low Mg^2+^/37 °C and 9.7% MeOH condition, the R1-09/R1-33 combination also performed about 1.4-times better than WT sequences in sfGFP expression (**Suppl. Table 1**).

**Figure 4:**
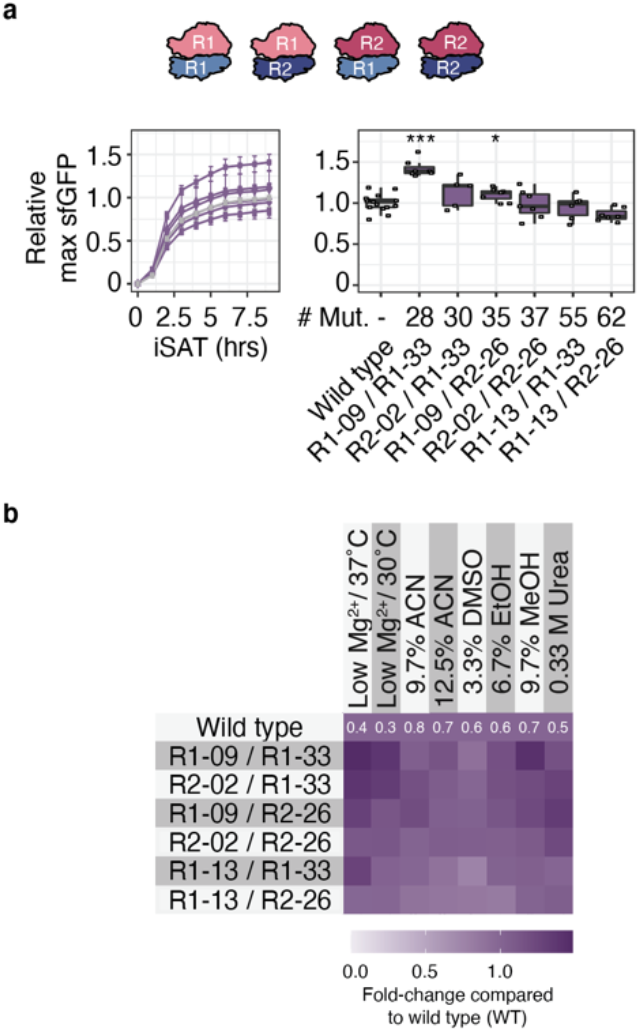
Eterna ribosomes confer beneficial phenotypes in vitro. Two 23S rRNA designs and three 16S rRNA designs with tolerant iSAT stress phenotypes were combined and tested in iSAT under various conditions: (a) optimal conditions, or (b) stress conditions. sfGFP expression in iSAT was determined by fluorescence and normalized to max sfGFP of pT7-rrnB-wild type at optimal iSAT conditions. Fold-changes in (b) illustrate iSAT activities normalized to wild type performance under solvent condition. ACN: acetonitrile, DMSO: dimethylsulfoxide, EtOH: ethanol, MeOH: methanol. Data are shown as boxplots with error bars representing s.d. (a) or mean (b); n ≥ 3.

### Community science design ribosomes are functional in vivo

A key challenge of ribosome design is that many rRNA mutants are dominantly lethal. Despite this challenge, we wondered if the community science designed ribosomes could support life. To test this, we individually cloned the R2 and “Combined” designs into the pL-rrnB plasmid, expressing the *rrnB* operon from a temperature-sensitive promoter, pL^37^, and conferring carbenicillin resistance. These plasmids were then individually transformed into the *Escherichia coli* SQ171fg strain^12^, which was evolved from the SQ171 strain^38^. The SQ171fg strain lacks chromosomal rRNA alleles and lives on the pCSacB plasmid which carries an rRNA operon encoding a tethered ribosome, Ribo-T v2^39^ and the tRNA67 plasmid encoding missing tRNA genes. The pCSacB plasmid also contains a counter-selectable marker sacB gene which confers sucrose sensitivity and a kanamycin resistance cassette. Transformed SQ171fg cells were grown in the presence of carbenicillin and sucrose, individual colonies were picked and tested for loss of kanamycin resistance, indicating loss of plasmid pCSacB-RiboT v2, and resistance to carbenicillin, indicating the presence of the corresponding pL-rrnB designs. Loss of pCSacB-RiboT v2 and presence and accuracy of the pL-rrnB-R2 plasmids was verified by Sanger sequencing. Strikingly, 18 of 20 R2 16S rRNA and 17 of 20 R2 23S rRNA designs support life— of the four designs that are non-functional in iSAT, three were lethal *in vivo* (R2-22, R2-23, R2-36), and one design with only about 50% iSAT activity (R2-13) harboring a mutation in the central 16S rRNA pseudoknot was lethal as well (**Figure 5a, b**). Interestingly, one design is inactive in iSAT, but functional in cells (R2-25); suggesting it may suffer from assembly defects *in vitro* which can be compensated for *in vivo*.

**Figure 5:**
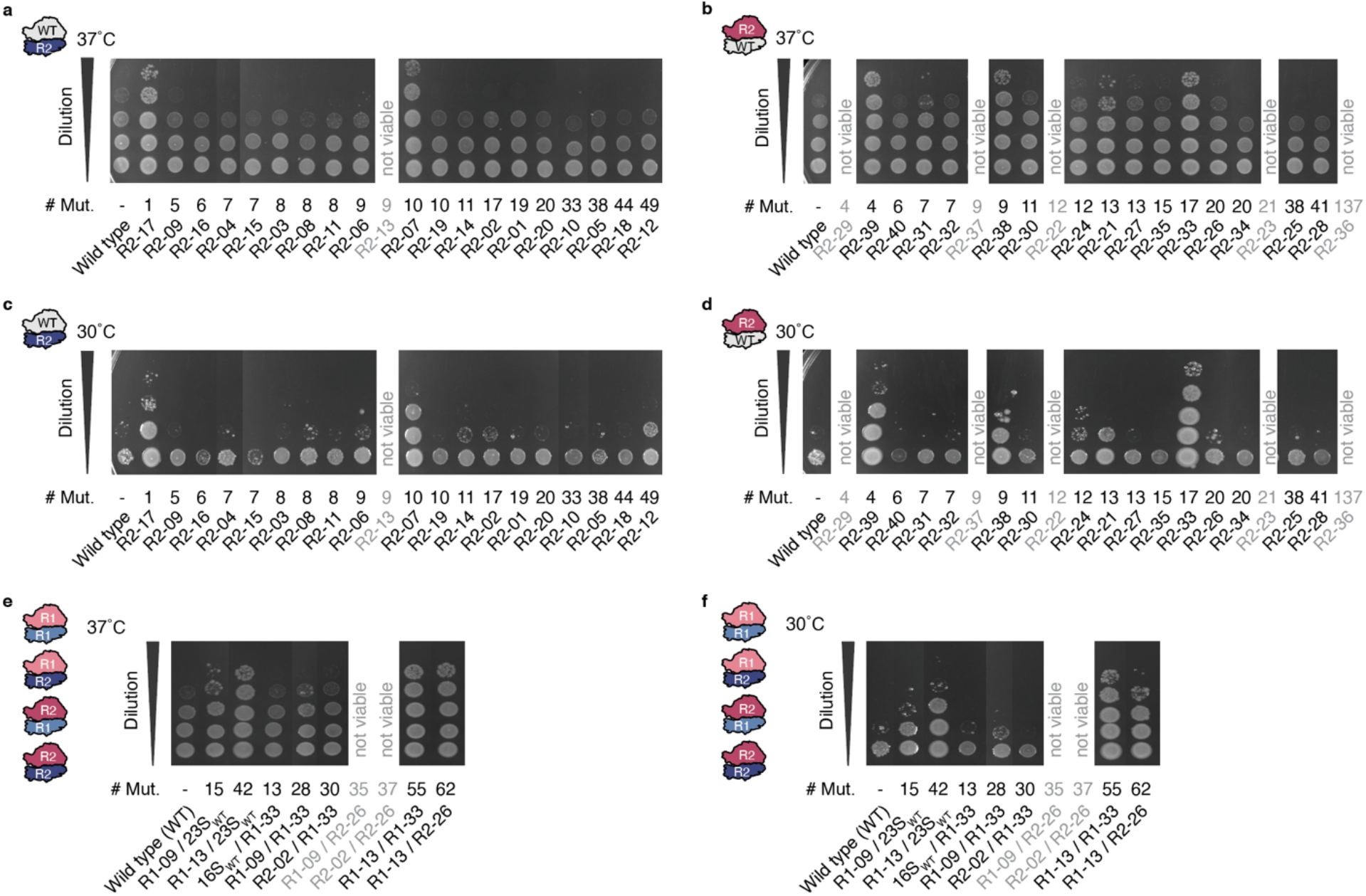
Eterna discovers diverse rRNAs which are functional *in vivo*. **a-f**) Spotted SQ171fg cells growing with pL-rrnB-wild type and pL-rrnB-R2 16S rRNA (**a**) and 23S rRNA (**b**) designs imaged after 24 hours at 37 °C or 72 hours at 30 °C. Stationary cells were diluted to an OD600 = 1, diluted stepwise 1:10, and spotted onto LB + Carb100 plates. Mut.: mutations, WT: wild type. Data representative of n=3 independent *in vivo* growth experiments.

We investigated how the ribosomes containing Eterna rRNA designs affect growth on solid nutritionally rich (LB) media. For this, we performed spot plating assays (**Figure 5a–d; Suppl. Figure 15**). Surprisingly, most of the R2 and combined 16S and 23S rRNA designs showed growth phenotypes on plates similar to WT, and 5 strains (R2-10, R2-21, R2-33, R2-38, R2-39) grew better than WT on plates (**Figure 5a, b; Suppl. Figure 15a, c**). When incubating the plates at sub-optimal temperature (30 °C) the phenotypes were even more pronounced and additional strains showed growth phenotypes (R2-12, R2-24); **Figure 5c, d; Suppl. Figure 15b, d**). We also assessed the functionality of the combinatorial design *in vivo* (**Figure 5e, f; Suppl. Figure 16**), and found that, except for two ribosomes designs (R1-09/R2-26 and R1-13/R2-26), the combined mutant ribosomes also support life. Surprisingly, two highly mutated ribosomes containing 55 and 62 mutations (combinatorial designs R1-13/R1-33 and R1-13/R2-26) show improved growth at 37 °C and 30 °C. This growth phenotype presumably arises from 16S rRNA design R1-13, which also confers improved growth when combined with WT 23S rRNA. Taken together, our results demonstrate the functionality of Eterna-designed ribosomes both *in vitro* and *in vivo*.

## DISCUSSION

In this study, we present an integrated computational and experimental workflow for constructing mutant ribosomes. This was accomplished by creating a DBTL pipeline that relies on game developers to create rRNA puzzles for stabilizing secondary structure, community scientists to design rRNA sequences, and lab scientists to carry out high-throughput, *in vitro* reactions to assess ribosome synthesis, assembly, and translation. We applied this pipeline to create mutant ribosomes that are functionally active.

Our work has several important features. First, we found that mutant rRNA sequences confer beneficial phenotypes. *In vitro*, we showed that ribosomes constructed and assessed in iSAT outperformed WT ribosomes in *in vitro* translation under standard conditions, and were even more tolerant in stress phenotypes (e.g., protein expression in organic solvents). *In vivo*, we were surprised to find that more than 85% of designed ribosomes tested could support life, with more than 10% demonstrating improved growth phenotypes in strains that live exclusively off community scientist designed ribosomes. Second, our data provided new insights into ribosome sequence-to-function relationships. For example, we found that mutations that slightly improve secondary structure energetics, or those that remove base repeats (R2-24), can be well tolerated, and that these effects can be amplified when combining mutations from well-performing designs (R2-02, R2-33). Surprisingly, we found that exchanging nucleotides against nucleotides from other gammaproteobacterial can improve protein translation *in vitro* (R1-13, R2-26) and growth *in vivo* (R1-13, R2-12, R2-38) at optimal and low temperature conditions. Third, we applied crowdsourcing to ribosome design for the first time. Crowdsourcing design presents complex problems to a diverse population of citizen scientists, resulting in a variety of solutions that, taken together, can avoid trapping by local optima in a fashion reminiscent of classic nonconvex optimization algorithms^40^. For example, in our case, we found that community scientists were able to improve at each stage of the process (from R0 through R2), working together to infer how to iterate on a family of solutions in light of newly disseminated experimental data.

Looking forward, our Eterna-based DBTL approach to crowdsourcing ribosome design may provide a rapid and powerful strategy for developing engineered ribosomes for synthetic and chemical biology. This could be used to deepen our understanding of the ribosome’s RNA-based active site, make simpler ribosomes to fill in knowledge gaps in the origins of life, and tailor the ribosome active site to accommodate novel monomers to yield new classes of enzymes, therapeutics, and materials.

## ACKNOWLEDGMENTS

This work was supported by the Army Research Office (W911NF-16-1-0372, M.C.J.), the David and Lucile Packard Foundation (to M.C.J.), the Camille Dreyfus Teacher-Scholar Program (to M.C.J.), the National Institutes of Health (R35 GM122579 to R.D.), and a Stanford Medicine Discovery Innovation Award (to R.D.). We thank Ashty Karim for helpful comments and discussions on the manuscript. We thank John Nicol for supporting the development of tools for the ribosome puzzles.

## AUTHOR CONTRIBUTIONS

A.K., A.D., C.K., and Y.K. conducted experiments. A.K. and A.M.W. analyzed data. A.K., A.M.W., R.D. and M.C.J. designed and interpreted experiments. Eterna puzzles and tools were designed and crowdsourced by A.M.W., R.D., R.W., J.R, J.A.L., E.F., J.T.. The manuscript was written by A.K., A.M.W., M.C.J. and R.D.

## DATA AND MATERIAL AVAILABILITY

rRNA designs are available from the authors upon request, and accessible on the Eterna website.

## METHODS

### S150 extract preparation

*Escherichia coli* MRE600 cells for S150 extract and TP70 preparation were grown in 2x YPTG at 37 °C until OD600 = 3.0. Cells were pelleted, washed three times in S150 lysis buffer (20 mM Tris-HCl (pH 7.2 at 4 °C), 100 mM NH_4_Cl, 10 mM MgCl_2_, 0.5 mM EDTA, 2 mM DTT), flash frozen in liquid nitrogen, and stored at −80 °C. For cell lysis, 4 g of cells were resuspended in S150 lysis buffer at a ratio of 5 ml of buffer per 1 g of cells and supplemented with 200 μl of Halt Protease Inhibitor Cocktail (Thermo Fisher Scientific Inc.) and 75 μl RNase Inhibitor (Qiagen) per 4 g of cells. The cells were lysed at ~ 20’000 psi with an EmulsiFlex-C3 homogenizer (Avestin). Lysate was supplemented with an equivalent dose of RNase Inhibitor and 3 μl of 1M DTT per ml suspension and clarified twice by centrifugation at 30’000 x g and 4 °C for 30 min. Resulting S30 extract was recovered and layered in a 1:1 volumetric ratio on a high sucrose cushion composed of Buffer B (20 mM Tris-HCl (pH 7.2 at 4 °C), 500 mM NH_4_Cl, 10 mM MgCl_2_, 0.5 mM EDTA, 2 mM DTT, 37.7% sucrose) into Ti70 ultracentrifuge tubes. Samples were centrifuged at 90,000 x g and 4 °C for 20 h. Clear ribosome pellets were used for TP70 preparation and supernatants recovered and spun at 150,000 x g and 4 °C for an additional 4 h. The top two-thirds of the supernatants were collected without disturbing the pellet and dialyzed in SnakeSkin dialysis tubing (Thermo Fisher Scientific; 3.5 kDa MWCO) against 50 volumes of high-salt S150 extract buffer (10 mM Tris-OAc (pH 7.5 at 4 °C), 10 mM Mg(OAc)_2_, 20 mM NH_4_OAc, 30 mM KOAc, 200 mM KGlu, 1 mM spermidine, 1 mM putrescine, 1 mM DTT). Four dialysis steps with fresh dialysis buffer were performed: three steps for 2 h and a final step overnight. Extracts were then clarified at 4’000 × g for 10 min and concentrated to ~4 mg/mL total protein concentration using 3 kDa molecular weight cutoff (MWCO) Centriprep concentrators (EMD Millipore) to account for dilution during preparation. S150 extract samples were aliquoted, flash-frozen, and stored at −80 °C. Protein concentration was determined using Bradford assay with bovine serum albumin (BSA; BioRad) as a standard.

### Total protein of 70S ribosomes (TP70) preparation

Clear ribosome pellets from S150 extract preparation were washed and resuspended in 10 mM Tris-OAc pH 7.5, 60 mM NH_4_Cl, 7.5 mM Mg(OAc)_2_, 0.5 mM EDTA, and 2 mM DTT. Concentration of resuspended ribosomes was determined from A260 NanoDrop readings (1 A260 unit of 70S = 24 pmol 70S^41^). Ribosomes were aliquoted, flash-frozen and stored at −80 °C until further use. To precipitate rRNA, two volumes of glacial acetic acid were added to purified 70S ribosomes in Buffer C (10 mM Tris-OAc (pH 7.5 at 4 °C), 60 mM NH_4_Cl, 7.5 mM Mg(OAc)_2_, 9.5 mM EDTA, 2 mM dithiothreitol (DTT)) with 0.2 mM spermine and 2 mM spermidine, and 100 mM Mg(OAc)_2_. Samples were mixed well and centrifuged at 16,000 × g for 30 min to pellet rRNA. Supernatants containing ribosomal proteins was collected, mixed with five volumes of chilled acetone and stored at −20 °C overnight. Precipitated protein was collected by centrifugation at 10’000 × g for 30 min, dried, and resuspended in simplified high-salt/ urea buffer (50 mM HEPES (pH 7.6 at RT), 10 mM Mg(Glu)_2_, 200 mM KGlu, 0.5 mM EDTA, 2 mM DTT, 1 mM putrescine, 1 mM spermidine, 6 M urea). Protein sample was then transferred to midi-size 1 kDa MWCO Tube-O-Dialyzers (G-Biosciences) and first dialyzed against 100 volumes of simplified high-salt buffer with urea overnight, then three times against 100 volumes of simplified high-salt buffer without urea for 90 min each. The dialyzed protein sample was next clarified at 4’000 × g for 10 min, and the supernatant’s protein concentration determined from A230 NanoDrop readings (1 A230 unit of TP70 = 240 pmol TP70 ^39^). TP70 samples were aliquoted, flash-frozen, and stored at −80 °C until use.

### iSAT reactions

iSAT reactions were set-up as previously described ^8,30,40,41^. Briefly, 6.5 μl reactions were prepared by mixing salts, substrates, cofactors and additives (7.5 mM Mg(Glu)_2_, 167 mM K(Glu), 1.2 mM ATP, 0.85 mM GTP, 0.85 mM UTP, 0.85 mM CTP, 0.034 mg/ml folinic acid, 0.1706 mg/ml tRNAs, 0.33 mM NAD, 0.27 mM CoA, 4 mM oxalic acid, 1 mM putrescine, 1.5 mM spermidine, 57 mM HEPES, 3 mM amino acids, 42 mM PEP, 4% PEG8000, 2 mM DTT), with 4 nM sfGFP reporter plasmid, 4 nM pT7rrnB construct, 60 μg/ml T7 RNA polymerase, 200 nM TP70, and S150 extract. Reactions were incubated in 384-well plates (Greiner, catalog number 781096) at 37 °C in a BioTek Synergy H1 plate reader, and fluorescence of superfolder GFP (sfGFP) was monitored (excitation: 450–490 nm, emission: 510–530 nm) over the course of the reaction.

### iSAT plasmid construction

DNA templates of 16S and 23S rRNA designed by community scientists (Eterna designs) were synthesized by Twist Biosciences and exchanged against wild type 16S rRNA and 23S rRNA of the 7311-bp plasmid pT7rrnB carrying the Escherichia coli *rrnB* operon under the control of the T7 promoter and the β-lactamase resistance gene as a selective marker (Supplemental data – Plasmid sequences).

### Plasmid construction for *in vivo* tests

DNA templates of 16S and 23S rRNA round 2 Eterna designs were synthesized by Twist Biosciences and exchanged against wild type 16S rRNA or mutant 23S rRNA A2058G of the 7415-bp plasmid pLrrnB carrying the *Escherichia coli rrnB* operon under the control of the pL-G-12T promoter^43^ and the β-lactamase resistance gene as a selective marker (Supplemental data – Plasmid sequences). Constructs carrying the 16S rRNA Eterna designs therefore harbored the A2058G point mutation in the 23S. This point mutation was corrected back to wild type by site-directed mutagenesis using Q5® Hot Start High-Fidelity DNA Polymerase (New England Biolabs) and primers: ACGGAAAGACCCCGTGAACC and CTTGCCGCGGGTACACTGC. Linear PCR products were purified using DNA Clean and Concentrator-5 kit (Zymo Research), DpnI digested, phosphorylated by T4 PNK (New England Biolabs), blunt-end ligated using T4 ligase (New England Biolabs), transformed into 50 μL of electrocompetent POP cells, recovered in 800 μL SOC media, plated onto LB-agar/ carbenicillin plates and grown at 30 °C. Clones were picked, streaked out onto fresh LB-agar/ carbenicillin plates, and grown over night in 3 ml of LB / carbenicillin media at 30 °C. Plasmids were prepped (Zymo Research) and rrnB operons sequence-verified using sanger sequencing (Northwestern University Sanger Sequencing Facility).

### Replacement of Ribo-Tv2 by Eterna-pLrrnB plasmid in SQ171fg cells

SQ171fg cells harboring the Ribo-Tv2 plasmid containing the kanamycin resistance gene as selection marker were transformed with pLrrnB plasmids carrying the 16S and 23S rRNA Eterna designs and the β-lactamase resistance gene. In brief, 20-100 ng of an Eterna-pLrrnB plasmid was transformed into 50 μL of electrocompetent cells. Cells were resuspended in 850 μL of SOC media and incubated for 1 h at 37 °C with shaking. 250 μL of recovering cells were transferred to 1.75 ml of SOC containing 50 μg/ ml of carbenicillin and 0.25% sucrose (final concentrations) and grown for 16-18 hours at 37 °C with shaking. Cells were pelleted and plated on LB-agar plates containing 50 μg/ ml carbenicillin and 5% sucrose. Colonies were tested for loss of the original Ribo-Tv2 plasmid and containment of the desired pLrrnB plasmid by selecting clones only living on LB-agar/ carbenicillin plates, but not on LB-agar/ kanamycin plates. The presence and correctness of the Eterna-design rrnB operon in identified clones was sanger sequence-verified.

### Spotting assay

Eterna-pLrrnB-containing SQ171fg cells were grown overnight in 3 mL of LB media containing 75 μg/ ml of carbenicillin. The cultures were diluted to OD600 = 1, 0.1, 0.01, 0.001, and 0.0001 with water. If not stated otherwise, 3 μl of the dilutions were spotted onto LB-agar/ 100 μg/ ml carbenicillin plates and grown for 24 hours at 37 °C or 72 hours at 30 °C.

### Computational prediction (CP) control

In order to provide a realistic comparison for the designs from the Eterna pilot round, a Python script was used to launch hundreds of Metropolis criterion Monte Carlo trajectories starting from the wild type sequence for the 16S and 23S rRNAs. These trajectories attempted to make mutations to the rRNA that would minimize the difference in energy between the “delta” – the energy of the sequence’s minimum free energy structure in the Vienna2 engine and the energy of the “target” structure used in the corresponding pilot round puzzle. Moves reducing “delta” were accepted unconditionally, while moves increasing “delta” were accepted conditionally (if they passed the Metropolis criterion). Following these trajectories, the resulting set of sequences were taken together and for each mutation count found in an Eterna design, the CP sequence with the lowest “delta” was selected. Scripts to carry out these simulations are included at https://github.com/everyday847/vienna_guided_mc. 8544 sequences were generated for the 23S rRNA, and 40689 sequences were generated for the 16S rRNA.

### Design of ribosome puzzles

The initial pilot round ribosome puzzles were created with the conventional tools already available in Eterna, which permitted the specification of a sequence and secondary structure, an energy model, and some design restrictions (i.e., that certain bases could not be mutated (i.e., “locked”) and that at most a certain number of mutations would be tolerated).

For that round, we prompted players to minimize the energy gap for each rRNA sequence between its experimental secondary structure and its minimal free energy (MFE) secondary structure. So that players could receive real-time feedback on their design progress, we created 16S and 23S rRNA puzzles and equipped them with the LinearFold implementation of the Vienna2 energy model^42^. As a precaution, we limited the total number of mutations to 5% of the 16S and 23S rRNA (76 and 145 mutations, respectively). We additionally prohibited 40 nt in the 16S rRNA and 70 nt in the 23S rRNA from being mutated; these “locked” bases included contacts with ribosomal proteins, tertiary contacts within one ribosomal subunit or across both subunits, or they formed base pairs that could not be recovered in the Vienna energy model (e.g., pseudoknots or single base pairs bridging two individually unstable two-way junctions (see Supplemental Data).

Eterna developers created several new features for the first full round of puzzles. To ameliorate the innumerable overlaps resulting from the naïve layout algorithm available in Eterna, the puzzles were modified to accept a “custom” layout for the target structure, whereby the correct RNA secondary structure had fixed Cartesian coordinates for each nucleotide, and every time a junction’s orientation was solved in “natural mode”, it would “snap” to the target mode coordinates, eliminating all target mode overlaps and improving the situation in natural mode. Furthermore, to provide more detailed feedback to players, a new constraint was implemented using sequence conservation of each nucleotide in the rRNA sequences across gammaproteobacteria. This “IUPAC constraint,” named for how sequence variation was encoded, ensured that players had real-time access to the mutations that self-evidently are tolerated in related organisms, thus allowing players to vote for solutions that might have many mutations, but relatively few “IUPAC violations.” The first round also included several “subpuzzles” permitting players to explore the individual domains from these large molecules in more detail.

For the second round, two significant changes were made to improve the 16S rRNA puzzle specifically. First, because the energy models used in Eterna – especially for ribosome puzzles – omit pseudoknots, the 16S helix ‘h2’, which is properly a pseudoknot, had only been modeled implicitly via IUPAC constraints or locks. In the second round, a second version of the 16S rRNA puzzle was provided with h2 defined instead of h1, permitting players to test their designs in each structural context. Second, the sequence used for the pilot and R1 16S rRNA was only 1534 nucleotides, omitting the aSD sequence. In the R2 16S rRNA puzzles, the aSD sequence appears and is locked to its WT sequence identity. The science team also created new resources that the players could use as a supplement to their design process. Several players concerned with the prospect of disrupting contacts with rProteins asked the science team for annotations of what nucleotides contacted proteins in each rRNA. The science team used a 3D structure of the *E. coli* ribosome (PDB code: 4YBB^35^) to annotate each protein-contacting residue and, importantly, to annotate which contacts directly influenced the allowed nucleotides at that position (i.e., a contact through the phosphate backbone does not constrain the nucleotide sequence, a contact with adenosine N7 could be satisfied by guanosine, but a contact with guanosine’s keto group could not be satisfied by another nucleotide).

