## Supplemental Data for "Community science designed ribosomes with beneficial phenotypes"

### Contents

1. Supplemental details on ribosome puzzle definitions
2. Plasmid sequences
3. Supplemental Figures

In addition to the following supplemental details, sequences, and figures, please find a Suppl. Table (Excel spreadsheet) containing all designed rRNAs' parameters and experimental data as well as diagrams of each of the rRNAs designed in this study supplied as a single .zip archive.

### 1. Supplemental details on ribosome puzzle definitions

*Energetic rationale for base locks*

Most base locks were chosen based on intra- or inter-subunit tertiary contacts, particularly when Watson-Crick, and protein-RNA contacts that would both enormously influence the folding energetics of the rRNA under investigation but could not be represented in a secondary structure folding model. Pseudoknotted residues were also locked; folding the ribosome with a pseudoknot-aware secondary structure model would be both physically unrealistic (ribosome folding is chaperoned; the only pseudoknots likely to form in designed ribosomes are those that form in the wild type) and computationally intractable.

Some "singlet" base pairs, however, were also locked. In large, folded RNAs, tertiary folding influences the secondary structure ensemble and can render stable features that otherwise might struggle to form. "Singlet" base pairs – those that do not form part of a secondary structure stem – are not favorable on their own, since they contribute no stabilizing stacking energy, and in energy models like Vienna will typically destabilize large loops.

Because this "secondary structure" constraint could not be satisfied in the energy model, we omitted it from the target secondary structure to make the objective more achievable for players, but we locked the nucleotides to ensure that these destabilized bases were not mutated.

### 2. Plasmid sequences

pL-rrnB-wild type:
GATCTCTCACCTACCAAACAATGCCCCCCTGCAAAAAATAAATTCATATAAAAAACATACAG
ATAACCATCTGCGGTGATAAATTATCTCTGGCGGTGTTGACATAAATACCACTGGCGGTTAT
ACTGAGCACGGGTACCGGCCGCTGAGAAAAAGCGAAGCGGCACTGCTCTTTAACAATTTA
TCAGACAATCTGTGTGGGCACTCGAAGATACGGATTCTTAACGTCGCAAGACGAAAAATGA
ATACCAAGTCTCAAGAGTGAACACGTAATTCATTACGAAGTTTAATTCTTTGAGCGTCAAAC
TTTTAAATTGAAGAGTTTGATCATGGCTCAGATTGAACGCTGGCGGCAGGCCTAACACATG
CAAGTCGAACGGTAACAGGAAGAAGCTTGCTTCTTTGCTGACGAGTGGCGGACGGGTGAG
TAATGTCTGGGAAACTGCCTGATGGAGGGGGATAACTACTGGAAACGGTAGCTAATACCG
CATAACGTCGCAAGACCAAAGAGGGGGACCTTCGGGCCTCTTGCCATCGGATGTGCCCAG
ATGGGATTAGCTAGTAGGTGGGGTAACGGCTCACCTAGGCGACGATCCCTAGCTGGTCTG
AGAGGATGACCAGCCACACTGGAACTGAGACACGGTCCAGACTCCTACGGGAGGCAGCA
GTGGGGAATATTGCACAATGGGCGCAAGCCTGATGCAGCCATGCCGCGTGTATGAAGAAG
GCCTTCGGGTTGTAAAGTACTTTCAGCGGGGAGGAAGGGAGTAAAGTTAATACCTTTGCTC
ATTGACGTTACCCGCAGAAGAAGCACCGGCTAACTCCGTGCCAGCAGCCGCGGTAATACG
GAGGGTGCAAGCGTTAATCGGAATTACTGGGCGTAAAGCGCACGCAGGCGGTTTGTTAAG
TCAGATGTGAAATCCCCGGGCTCAACCTGGGAACTGCATCTGATACTGGCAAGCTTGAGT
CTCGTAGAGGGGGGTAGAATTCCAGGTGTAGCGGTGAAATGCGTAGAGATCTGGAGGAAT
ACCGGTGGCGAAGGCGGCCCCCTGGACGAAGACTGACGCTCAGGTGCGAAAGCGTGGG
GAGCAAACAGGATTAGATACCCTGGTAGTCCACGCCGTAAACGATGTCGACTTGGAGGTT
GTGCCCTTGAGGCGTGGCTTCCGGAGCTAACGCGTTAAGTCGACCGCCTGGGGAGTACG
GCCGCAAGGTTAAAACTCAAATGAATTGACGGGGGCCCGCACAAGCGGTGGAGCATGTG
GTTTAATTCGATGCAACGCGAAGAACCTTACCTGGTCTTGACATCCACGGAAGTTTTCAGA
GATGAGAATGTGCCTTCGGGAACCGTGAGACAGGTGCTGCATGGCTGTCGTCAGCTCGTG
TTGTGAAATGTTGGGTTAAGTCCCGCAACGAGCGCAACCCTTATCCTTTGTTGCCAGCGGT
CCGGCCGGGAACTCAAAGGAGACTGCCAGTGATAAACTGGAGGAAGGTGGGGATGACGT

CAAGTCATCATGGCCCTTACGACCAGGGCTACACACGTGCTACAATGGCGCATACAAAGA
GAAGCGACCTCGCGAGAGCAAGCGGACCTCATAAAGTGCGTCGTAGTCCGGATTGGAGTC
TGCAACTCGACTCCATGAAGTCGGAATCGCTAGTAATCGTGGATCAGAATGCCACGGTGAA
TACGTTCCCGGGCCTTGTACACACCGCCCGTCACACCATGGGAGTGGGTTGCAAAAGAAG
TAGGTAGCTTAACCTTCGGGAGGGCGCTTACCACTTTGTGATTCATGACTGGGGTGAAGTC
GTAACAAGGTAACCGTAGGGGAACCTGCGGTTGGATCACCTCCTTACCTTAAAGAAGCGTA
CTTTGTAGTGCTCACACAGATTGTCTGATAGAAAGTGAAAAGCAAGGCGTTTACGCGTTGG
GAGTGAGGCTGAAGAGAATAAGGCCGTTCGCTTTCTATTAATGAAAGCTCACCCTACACGA
AAATATCACGCAACGCGTGATAAGCAATTTTCGTGTCCCCTTCGTCTAGAGGCCCAGGACA
CCGCCCTTTCACGGCGGTAACAGGGGTTCGAATCCCCTAGGGGACGCCACTTGCTGGTTT
GTGAGTGAAAGTCGCCGACCTTAATATCTCAAAACTCATCTTCGGGTGATGTTTGAGATATT
TGCTCTTTAAAAATCTGGATCAAGCTGAAAATTGAAACACTGAACAACGAGAGTTGTTCGTG
AGTCTCTCAAATTTTCGCAACACGATGATGAATCGAAAGAAACATCTTCGGGTTGTGAGGTT
AAGCGACTAAGCGTACACGGTGGATGCCCTGGCAGTCAGAGGCGATGAAGGACGTGCTA
ATCTGCGATAAGCGTCGGTAAGGTGATATGAACCGTTATAACCGGCGATTTCCGAATGGG
GAAACCCAGTGTGTTTCGACACACTATCATTAACTGAATCCATAGGTTAATGAGGCGAACC
GGGGGAACTGAAACATCTAAGTACCCCGAGGAAAAGAAATCAACCGAGATTCCCCCAGTA
GCGGCGAGCGAACGGGGAGCAGCCCAGAGCCTGAATCAGTGTGTGTGTTAGTGGAAGCG
TCTGGAAAGGCGCGCGATACAGGGTGACAGCCCCGTACACAAAAATGCACATGCTGTGAG
CTCGATGAGTAGGGCGGGACACGTGGTATCCTGTCTGAATATGGGGGGACCATCCTCCAA
GGCTAAATACTCCTGACTGACCGATAGTGAACCAGTACCGTGAGGGAAAGGCGAAAAGAA
CCCCGGCGAGGGGAGTGAAAAAGAACCTGAAACCGTGTACGTACAAGCAGTGGGAGCAC
GCTTAGGCGTGTGACTGCGTACCTTTTGTATAATGGGTCAGCGACTTATATTCTGTAGCAA
GGTTAACCGAATAGGGGAGCCGAAGGGAAACCGAGTCTTAACTGGGCGTTAAGTTGCAGG
GTATAGACCCGAAACCCGGTGATCTAGCCATGGGCAGGTTGAAGGTTGGGTAACACTAAC
TGGAGGACCGAACCGACTAATGTTGAAAAATTAGCGGATGACTTGTGGCTGGGGGTGAAA
GGCCAATCAAACCGGGAGATAGCTGGTTCTCCCCGAAAGCTATTTAGGTAGCGCCTCGTG
AATTCATCTCCGGGGGTAGAGCACTGTTTCGGCAAGGGGGTCATCCCGACTTACCAACCC
GATGCAAACTGCGAATACCGGAGAATGTTATCACGGGAGACACACGGCGGGTGCTAACGT
CCGTCGTGAAGAGGGAAACAACCCAGACCGCCAGCTAAGGTCCCAAAGTCATGGTTAAGT
GGGAAACGATGTGGGAAGGCCCAGACAGCCAGGATGTTGGCTTAGAAGCAGCCATCATTT
AAAGAAAGCGTAATAGCTCACTGGTCGAGTCGGCCTGCGCGGAAGATGTAACGGGGCTAA
ACCATGCACCGAAGCTGCGGCAGCGACGCTTATGCGTTGTTGGGTAGGGGAGCGTTCTGT
AAGCCTGCGAAGGTGTGCTGTGAGGCATGCTGGAGGTATCAGAAGTGCGAATGCTGACAT
AAGTAACGATAAAGCGGGTGAAAAGCCCGCTCGCCGGAAGACCAAGGGTTCCTGTCCAAC
GTTAATCGGGGCAGGGTGAGTCGACCCCTAAGGCGAGGCCGAAAGGCGTAGTCGATGGG
AAACAGGTTAATATTCCTGTACTTGGTGTTACTGCGAAGGGGGGACGGAGAAGGCTATGTT
GGCCGGGCGACGGTTGTCCCGGTTTAAGCGTGTAGGCTGGTTTTCCAGGCAAATCCGGAA
AATCAAGGCTGAGGCGTGATGACGAGGCACTACGGTGCTGAAGCAACAAATGCCCTGCTT
CCAGGAAAAGCCTCTAAGCATCAGGTAACATCAAATCGTACCCCAAACCGACACAGGTGGT
CAGGTAGAGAATACCAAGGCGCTTGAGAGAACTCGGGTGAAGGAACTAGGCAAAATGGTG
CCGTAACTTCGGGAGAAGGCACGCTGATATGTAGGTGAGGTCCCTCGCGGATGGAGCTGA
AATCAGTCGAAGATACCAGCTGGCTGCAACTGTTTATTAAAAACACAGCACTGTGCAAACA
CGAAAGTGGACGTATACGGTGTGACGCCTGCCCGGTGCCGGAAGGTTAATTGATGGGGTT
AGCGCAAGCGAAGCTCTTGATCGAAGCCCCGGTAAACGGCGGCCGTAACTATAACGGTCC
TAAGGTAGCGAAATTCCTTGTCGGGTAAGTTCCGACCTGCACGAATGGCGTAATGATGGC
CAGGCTGTCTCCACCCGAGACTCAGTGAAATTGAACTCGCTGTGAAGATGCAGTGTACCC
GCGGCAAGACGGAAAGACCCCGTGAACCTTTACTATAGCTTGACACTGAACATTGAGCCTT
GATGTGTAGGATAGGTGGGAGGCTTTGAAGTGTGGACGCCAGTCTGCATGGAGCCGACCT
TGAAATACCACCCTTTAATGTTTGATGTTCTAACGTTGACCCGTAATCCGGGTTGCGGACA
GTGTCTGGTGGGTAGTTTGACTGGGGCGGTCTCCTCCTAAAGAGTAACGGAGGAGCACGA

AGGTTGGCTAATCCTGGTCGGACATCAGGAGGTTAGTGCAATGGCATAAGCCAGCTTGAC
TGCGAGCGTGACGGCGCGAGCAGGTGCGAAAGCAGGTCATAGTGATCCGGTGGTTCTGA
ATGGAAGGGCCATCGCTCAACGGATAAAAGGTACTCCGGGGATAACAGGCTGATACCGCC
CAAGAGTTCATATCGACGGCGGTGTTTGGCACCTCGATGTCGGCTCATCACATCCTGGGG
CTGAAGTAGGTCCCAAGGGTATGGCTGTTCGCCATTTAAAGTGGTACGCGAGCTGGGTTT
AGAACGTCGTGAGACAGTTCGGTCCCTATCTGCCGTGGGCGCTGGAGAACTGAGGGGGG
CTGCTCCTAGTACGAGAGGACCGGAGTGGACGCATCACTGGTGTTCGGGTTGTCATGCCA
ATGGCACTGCCCGGTAGCTAAATGCGGAAGAGATAAGTGCTGAAAGCATCTAAGCACGAA
ACTTGCCCCGAGATGAGTTCTCCCTGACCCTTTAAGGGTCCTGAAGGAACGTTGAAGACG
ACGACGTTGATAGGCCGGGTGTGTAAGCGCAGCGATGCGTTGAGCTAACCGGTACTAATG
AACCGTGAGGCTTAACCTTACAACGCCGAAGCTGTTTTGGCGGATGAGAGAAGATTTTCAG
CCTGATACAGATTAAATCAGAACGCAGAAGCGGTCTGATAAAACAGAATTTGCCTGGCGGC
AGTAGCGCGGTGGTCCCACCTGACCCCATGCCGAACTCAGAAGTGAAACGCCGTAGCGC
CGATGGTAGTGTGGGGTCTCCCCATGCGAGAGTAGGGAACTGCCAGGCATCAAATAAAAC
GAAAGGCTCAGTCGAAAGACTGGGCCTTTCGTTTTATCTGTTGTTTGTCGGTGAACGCTCT
CCTGAGTAGGACAAATCCGCCGGGAGCGGATTTGAACGTTGCGAAGCAACGGCCCGGAG
GGTGGCGGGCAGGACGCCCGCCATAAACTGCCAGGCATCAAATTAAGCAGAAGGCCATC
CTGACGGATGGCCTTTTTGCGTTTCTACAAACTCTTCCTGTCGTCATATCTACAAGCCGGC
GCGCCGGGAAATGTGCGCGGAACCCCTATTTGTTTATTTTTCTAAATACATTCAAATATGTA
TCCGCTCATGAGACAATAACCCTGATAAATGCTTCAATAATATTGAAAAAGGAAGAGTATGA
GTATTCAACATTTCCGTGTCGCCCTTATTCCCTTTTTTGCGGCATTTTGCCTTCCTGTTTTTG
CTCACCCAGAAACGCTGGTGAAAGTAAAAGATGCTGAAGATCAGTTGGGTGCACGAGTGG
GTTACATCGAACTGGATCTCAACAGCGGTAAGATCCTTGAGAGTTTTCGCCCCGAAGAACG
TTTTCCAATGATGAGCACTTTTAAAGTTCTGCTATGTGGCGCGGTATTATCCCGTGTTGACG
CCGGGCAAGAGCAACTCGGTCGCCGCATACACTATTCTCAGAATGACTTGGTTGAGTACTC
ACCAGTCACAGAAAAGCATCTTACGGATGGCATGACAGTAAGAGAATTATGCAGTGCTGCA
ATAACCATGAGTGATAACACTGCGGCCAACTTACTTCTGACAACGATCGGAGGACCGAAG
GAGCTAACCGCTTTTTTGCACAACATGGGGGATCATGTAACTCGCCTTGATCGTTGGGAAC
CGGAGCTGAATGAAGCCATACCAAACGACGAGCGTGACACCACGATGCCTGCAGCAATGG
CAACAACGTTGCGCAAACTATTAACTGGCGAACTACTTACTCTAGCTTCCCGGCAACAATTA
ATAGACTGGATGGAGGCGGATAAAGTTGCAGGACCACTTCTGCGCTCGGCCCTTCCGGCT
AGCTGGTTTATTGCTGATAAATCTGGAGCCGGTGAGCGTGGGTCTCGCGGTATCATTGCA
GCACTGGGGCCAGATGGTAAGCCCTCCCGTATCGTAGTTATCTACACGACGGGGAGTCAG
GCAACTATGGATGAACGAAATAGACAGATCGCTGAGATAGGTGCCTCACTGATTAAGCATT
GGTAACTGCAGACCAAGTTTACTCATATATACTTTAGATTGATTTAAAACTTCATTTTTAATTT
AAAAGGATCTAGGTGAAGATCCTTTTTGATAATCTCATGACCAAAATCCCTTAACGTGAGTT
TTCGTTCCACTGAGCGTCAGACCCCGTAGAAAAGATCAAAGGATCTTCTTGAGATCCTTTTT
TTCTGCGCGTAATCTGCTGCTTGCAAACAAAAAAACCACCGCTACCAGCGGTGGTTTGTTT
GCCGGATCAAGAGCTACCAACTCTTTTTCCGAAGGTAACTGGCTTCAGCAGAGCGCAGATA
CCAAATACTGTCCTTCTAGTGTAGCCGTAGTTAGGCCACCACTTCAAGAACTCTGTAGCAC
CGCCTACATACCTCGCTCTGCTAATCCTGTTACCAGTGGCTGCTGCCAGTGGCGATAAGTC
GTGTCTTACCGGGTTGGACTCAAGACGATAGTTACCGGATAAGGCGCAGCGGTCGGGCTG
AACGGGGGGTTCGTGCACACAGCCCAGCTTGGAGCGAACGACCTACACCGAACTGAGATA
CCTACAGCGTGAGCTATGAGAAAGCGCCACGCTTCCCGAAGGGAGAAAGGCGGACAGGT
ATCCGGTAAGCGGCAGGGTCGGAACAGGAGAGCGCACGAGGGAGCTTCCAGGGGGAAA
CGCCTGGTATCTTTATAGTCCTGTCGGGTTTCGCCACCTCTGACTTGAGCGTCGATTTTTG
TGATGCTCGTCAGGGGGGCGGAGCCTATGGAAAAACGCCAGCAACGCGGCCTTTTTACG
GTTCCTGGCCTTTTGCTGGGCGGCCGC

pT7-rrnB-wild type:
TAATACGACTCACTATAGGGGCCGCTGAGAAAAAGCGAAGCGGCACTGCTCTTTAACAATT
TATCAGACAATCTGTGTGGGCACTCGAAGATACGGATTCTTAACGTCGCAAGACGAAAAAT
GAATACCAAGTCTCAAGAGTGAACACGTAATTCATTACGAAGTTTAATTCTTTGAGCGTCAA
ACTTTTAAATTGAAGAGTTTGATCATGGCTCAGATTGAACGCTGGCGGCAGGCCTAACACA
TGCAAGTCGAACGGTAACAGGAAGAAGCTTGCTTCTTTGCTGACGAGTGGCGGACGGGTG
AGTAATGTCTGGGAAACTGCCTGATGGAGGGGGATAACTACTGGAAACGGTAGCTAATAC
CGCATAACGTCGCAAGACCAAAGAGGGGGACCTTCGGGCCTCTTGCCATCGGATGTGCCC
AGATGGGATTAGCTAGTAGGTGGGGTAACGGCTCACCTAGGCGACGATCCCTAGCTGGTC
TGAGAGGATGACCAGCCACACTGGAACTGAGACACGGTCCAGACTCCTACGGGAGGCAG
CAGTGGGGAATATTGCACAATGGGCGCAAGCCTGATGCAGCCATGCCGCGTGTATGAAGA
AGGCCTTCGGGTTGTAAAGTACTTTCAGCGGGGAGGAAGGGAGTAAAGTTAATACCTTTGC
TCATTGACGTTACCCGCAGAAGAAGCACCGGCTAACTCCGTGCCAGCAGCCGCGGTAATA
CGGAGGGTGCAAGCGTTAATCGGAATTACTGGGCGTAAAGCGCACGCAGGCGGTTTGTTA
AGTCAGATGTGAAATCCCCGGGCTCAACCTGGGAACTGCATCTGATACTGGCAAGCTTGA
GTCTCGTAGAGGGGGGTAGAATTCCAGGTGTAGCGGTGAAATGCGTAGAGATCTGGAGGA
ATACCGGTGGCGAAGGCGGCCCCCTGGACGAAGACTGACGCTCAGGTGCGAAAGCGTGG
GGAGCAAACAGGATTAGATACCCTGGTAGTCCACGCCGTAAACGATGTCGACTTGGAGGT
TGTGCCCTTGAGGCGTGGCTTCCGGAGCTAACGCGTTAAGTCGACCGCCTGGGGAGTAC
GGCCGCAAGGTTAAAACTCAAATGAATTGACGGGGGCCCGCACAAGCGGTGGAGCATGT
GGTTTAATTCGATGCAACGCGAAGAACCTTACCTGGTCTTGACATCCACGGAAGTTTTCAG
AGATGAGAATGTGCCTTCGGGAACCGTGAGACAGGTGCTGCATGGCTGTCGTCAGCTCGT
GTTGTGAAATGTTGGGTTAAGTCCCGCAACGAGCGCAACCCTTATCCTTTGTTGCCAGCGG
TCCGGCCGGGAACTCAAAGGAGACTGCCAGTGATAAACTGGAGGAAGGTGGGGATGACG
TCAAGTCATCATGGCCCTTACGACCAGGGCTACACACGTGCTACAATGGCGCATACAAAGA
GAAGCGACCTCGCGAGAGCAAGCGGACCTCATAAAGTGCGTCGTAGTCCGGATTGGAGTC
TGCAACTCGACTCCATGAAGTCGGAATCGCTAGTAATCGTGGATCAGAATGCCACGGTGAA
TACGTTCCCGGGCCTTGTACACACCGCCCGTCACACCATGGGAGTGGGTTGCAAAAGAAG
TAGGTAGCTTAACCTTCGGGAGGGCGCTTACCACTTTGTGATTCATGACTGGGGTGAAGTC
GTAACAAGGTAACCGTAGGGGAACCTGCGGTTGGATCACCTCCTTACCTTAAAGAAGCGTA
CTTTGTAGTGCTCACACAGATTGTCTGATAGAAAGTGAAAAGCAAGGCGTTTACGCGTTGG
GAGTGAGGCTGAAGAGAATAAGGCCGTTCGCTTTCTATTAATGAAAGCTCACCCTACACGA
AAATATCACGCAACGCGTGATAAGCAATTTTCGTGTCCCCTTCGTCTAGAGGCCCAGGACA
CCGCCCTTTCACGGCGGTAACAGGGGTTCGAATCCCCTAGGGGACGCCACTTGCTGGTTT
GTGAGTGAAAGTCGCCGACCTTAATATCTCAAAACTCATCTTCGGGTGATGTTTGAGATATT
TGCTCTTTAAAAATCTGGATCAAGCTGAAAATTGAAACACTGAACAACGAGAGTTGTTCGTG
AGTCTCTCAAATTTTCGCAACACGATGATGAATCGAAAGAAACATCTTCGGGTTGTGAGGTT
AAGCGACTAAGCGTACACGGTGGATGCCCTGGCAGTCAGAGGCGATGAAGGACGTGCTA
ATCTGCGATAAGCGTCGGTAAGGTGATATGAACCGTTATAACCGGCGATTTCCGAATGGG
GAAACCCAGTGTGTTTCGACACACTATCATTAACTGAATCCATAGGTTAATGAGGCGAACC
GGGGGAACTGAAACATCTAAGTACCCCGAGGAAAAGAAATCAACCGAGATTCCCCCAGTA
GCGGCGAGCGAACGGGGAGCAGCCCAGAGCCTGAATCAGTGTGTGTGTTAGTGGAAGCG
TCTGGAAAGGCGCGCGATACAGGGTGACAGCCCCGTACACAAAAATGCACATGCTGTGAG
CTCGATGAGTAGGGCGGGACACGTGGTATCCTGTCTGAATATGGGGGGACCATCCTCCAA
GGCTAAATACTCCTGACTGACCGATAGTGAACCAGTACCGTGAGGGAAAGGCGAAAAGAA
CCCCGGCGAGGGGAGTGAAAAAGAACCTGAAACCGTGTACGTACAAGCAGTGGGAGCAC
GCTTAGGCGTGTGACTGCGTACCTTTTGTATAATGGGTCAGCGACTTATATTCTGTAGCAA
GGTTAACCGAATAGGGGAGCCGAAGGGAAACCGAGTCTTAACTGGGCGTTAAGTTGCAGG
GTATAGACCCGAAACCCGGTGATCTAGCCATGGGCAGGTTGAAGGTTGGGTAACACTAAC
TGGAGGACCGAACCGACTAATGTTGAAAAATTAGCGGATGACTTGTGGCTGGGGGTGAAA
GGCCAATCAAACCGGGAGATAGCTGGTTCTCCCCGAAAGCTATTTAGGTAGCGCCTCGTG

AATTCATCTCCGGGGGTAGAGCACTGTTTCGGCAAGGGGGTCATCCCGACTTACCAACCC
GATGCAAACTGCGAATACCGGAGAATGTTATCACGGGAGACACACGGCGGGTGCTAACGT
CCGTCGTGAAGAGGGAAACAACCCAGACCGCCAGCTAAGGTCCCAAAGTCATGGTTAAGT
GGGAAACGATGTGGGAAGGCCCAGACAGCCAGGATGTTGGCTTAGAAGCAGCCATCATTT
AAAGAAAGCGTAATAGCTCACTGGTCGAGTCGGCCTGCGCGGAAGATGTAACGGGGCTAA
ACCATGCACCGAAGCTGCGGCAGCGACGCTTATGCGTTGTTGGGTAGGGGAGCGTTCTGT
AAGCCTGCGAAGGTGTGCTGTGAGGCATGCTGGAGGTATCAGAAGTGCGAATGCTGACAT
AAGTAACGATAAAGCGGGTGAAAAGCCCGCTCGCCGGAAGACCAAGGGTTCCTGTCCAAC
GTTAATCGGGGCAGGGTGAGTCGACCCCTAAGGCGAGGCCGAAAGGCGTAGTCGATGGG
AAACAGGTTAATATTCCTGTACTTGGTGTTACTGCGAAGGGGGGACGGAGAAGGCTATGTT
GGCCGGGCGACGGTTGTCCCGGTTTAAGCGTGTAGGCTGGTTTTCCAGGCAAATCCGGAA
AATCAAGGCTGAGGCGTGATGACGAGGCACTACGGTGCTGAAGCAACAAATGCCCTGCTT
CCAGGAAAAGCCTCTAAGCATCAGGTAACATCAAATCGTACCCCAAACCGACACAGGTGGT
CAGGTAGAGAATACCAAGGCGCTTGAGAGAACTCGGGTGAAGGAACTAGGCAAAATGGTG
CCGTAACTTCGGGAGAAGGCACGCTGATATGTAGGTGAGGTCCCTCGCGGATGGAGCTGA
AATCAGTCGAAGATACCAGCTGGCTGCAACTGTTTATTAAAAACACAGCACTGTGCAAACA
CGAAAGTGGACGTATACGGTGTGACGCCTGCCCGGTGCCGGAAGGTTAATTGATGGGGTT
AGCGCAAGCGAAGCTCTTGATCGAAGCCCCGGTAAACGGCGGCCGTAACTATAACGGTCC
TAAGGTAGCGAAATTCCTTGTCGGGTAAGTTCCGACCTGCACGAATGGCGTAATGATGGC
CAGGCTGTCTCCACCCGAGACTCAGTGAAATTGAACTCGCTGTGAAGATGCAGTGTACCC
GCGGCAAGACGGAAAGACCCCGTGAACCTTTACTATAGCTTGACACTGAACATTGAGCCTT
GATGTGTAGGATAGGTGGGAGGCTTTGAAGTGTGGACGCCAGTCTGCATGGAGCCGACCT
TGAAATACCACCCTTTAATGTTTGATGTTCTAACGTTGACCCGTAATCCGGGTTGCGGACA
GTGTCTGGTGGGTAGTTTGACTGGGGCGGTCTCCTCCTAAAGAGTAACGGAGGAGCACGA
AGGTTGGCTAATCCTGGTCGGACATCAGGAGGTTAGTGCAATGGCATAAGCCAGCTTGAC
TGCGAGCGTGACGGCGCGAGCAGGTGCGAAAGCAGGTCATAGTGATCCGGTGGTTCTGA
ATGGAAGGGCCATCGCTCAACGGATAAAAGGTACTCCGGGGATAACAGGCTGATACCGCC
CAAGAGTTCATATCGACGGCGGTGTTTGGCACCTCGATGTCGGCTCATCACATCCTGGGG
CTGAAGTAGGTCCCAAGGGTATGGCTGTTCGCCATTTAAAGTGGTACGCGAGCTGGGTTT
AGAACGTCGTGAGACAGTTCGGTCCCTATCTGCCGTGGGCGCTGGAGAACTGAGGGGGG
CTGCTCCTAGTACGAGAGGACCGGAGTGGACGCATCACTGGTGTTCGGGTTGTCATGCCA
ATGGCACTGCCCGGTAGCTAAATGCGGAAGAGATAAGTGCTGAAAGCATCTAAGCACGAA
ACTTGCCCCGAGATGAGTTCTCCCTGACCCTTTAAGGGTCCTGAAGGAACGTTGAAGACG
ACGACGTTGATAGGCCGGGTGTGTAAGCGCAGCGATGCGTTGAGCTAACCGGTACTAATG
AACCGTGAGGCTTAACCTTACAACGCCGAAGCTGTTTTGGCGGATGAGAGAAGATTTTCAG
CCTGATACAGATTAAATCAGAACGCAGAAGCGGTCTGATAAAACAGAATTTGCCTGGCGGC
AGTAGCGCGGTGGTCCCACCTGACCCCATGCCGAACTCAGAAGTGAAACGCCGTAGCGC
CGATGGTAGTGTGGGGTCTCCCCATGCGAGAGTAGGGAACTGCCAGGCATCAAATAAAAC
GAAAGGCTCAGTCGAAAGACTGGGCCTTTCGTTTTATCTGTTGTTTGTCGGTGAACGCTCT
CCTGAGTAGGACAAATCCGCCGGGAGCGGATTTGAACGTTGCGAAGCAACGGCCCGGAG
GGTGGCGGGCAGGACGCCCGCCATAAACTGCCAGGCATCAAATTAAGCAGAAGGCCATC
CTGACGGATGGCCTTTTTGCGTTTCTACAAACTCTTCCTGTCGTCATATCTACAAGCCGGC
GCGCCAAATTGACAATTACTCATCCGGCTCGAATAATGTGTGGAACTTAAACACACACAGG
AGGAAAACATATGTCTATCCAGCACTTCCGTGTTGCGCTGATCCCGTTCTTCGCGGCGTTC
TGCCTGCCGGTTTTCGCGCACCCGGAAACCCTGGTTAAAGTTAAAGACGCGGAAGACCAG
CTGGGTGCGCGTGTTGGTTACATCGAACTGGACCTGAACTCTGGTAAAATCCTGGAATCTT
TCCGTCCGGAAGAACGTTTCCCGATGATGTCTACCTTCAAAGTTCTGCTGTGCGGTGCGGT
TCTGTCTCGTGTTGACGCGGGTCAGGAACAGCTGGGTCGTCGTATCCACTACTCTCAGAA
CGACCTGGTTGAATACTCTCCCGTTACCGAAAAACACCTGACCGACGGTATGACCGTTCGT
GAACTGTGCTCTGCGGCGATCACCATGTCTGACAACACCGCAGCGAACCTGCTGCTGACC
ACCATCGGTGGTCCGAAAGAACTGACCGCGTTCCTGCACAACATGGGCGACCACGTTACC

CGTCTGGACCGTTGGGAACCGGAACTGAACGAAGCGATCCCGAACGACGAACGTGACAC
CACCATGCCTGCGGCGATGGCGACCACCCTGCGTAAACTGCTGACCGGTGAACTGCTGAC
CCTGGCATCTCGTCAGCAGCTGATCGACTGGATGGAAGCGGACAAAGTTGCGGGTCCGCT
GCTGCGTTCTGCGCTGCCTGCGGGTTGGTTCATCGCGGACAAATCTGGTGCGGGTGAAC
GTGGTTCTCGTGGTATCATCGCGGCGCTGGGTCCGGACGGTAAACCGTCTCGTATCGTTG
TTATCTACACCACCGGTTCTCAGGCGACCATGGACGAACGTAACCGTCAGATCGCGGAAA
TCGGTGCGTCTCTGATTAAACACTGGTAAACTCACTCCTAGCCCGCCTAATAAGCGGGCTT
TTTTTCTGCAGACCAAGTTTACTCATATATACTTTAGATTGATTTAAAACTTCATTTTTAATTT
AAAAGGATCTAGGTGAAGATCCTTTTTGATAATCTCATGACCAAAATCCCTTAACGTGAGTT
TTCGTTCCACTGAGCGTCAGACCCCGTAGAAAAGATCAAAGGATCTTCTTGAGATCCTTTTT
TTCTGCGCGTAATCTGCTGCTTGCAAACAAAAAAACCACCGCTACCAGCGGTGGTTTGTTT
GCCGGATCAAGAGCTACCAACTCTTTTTCCGAAGGTAACTGGCTTCAGCAGAGCGCAGATA
CCAAATACTGTCCTTCTAGTGTAGCCGTAGTTAGGCCACCACTTCAAGAACTCTGTAGCAC
CGCCTACATACCTCGCTCTGCTAATCCTGTTACCAGTGGCTGCTGCCAGTGGCGATAAGTC
GTGTCTTACCGGGTTGGACTCAAGACGATAGTTACCGGATAAGGCGCAGCGGTCGGGCTG
AACGGGGGGTTCGTGCACACAGCCCAGCTTGGAGCGAACGACCTACACCGAACTGAGATA
CCTACAGCGTGAGCTATGAGAAAGCGCCACGCTTCCCGAAGGGAGAAAGGCGGACAGGT
ATCCGGTAAGCGGCAGGGTCGGAACAGGAGAGCGCACGAGGGAGCTTCCAGGGGGAAA
CGCCTGGTATCTTTATAGTCCTGTCGGGTTTCGCCACCTCTGACTTGAGCGTCGATTTTTG
TGATGCTCGTCAGGGGGGCGGAGCCTATGGAAAAACGCCAGCAACGCGGCCTTTTTACG
GTTCCTGGCCTTTTGCTGGT

### 3. Supplemental Figures

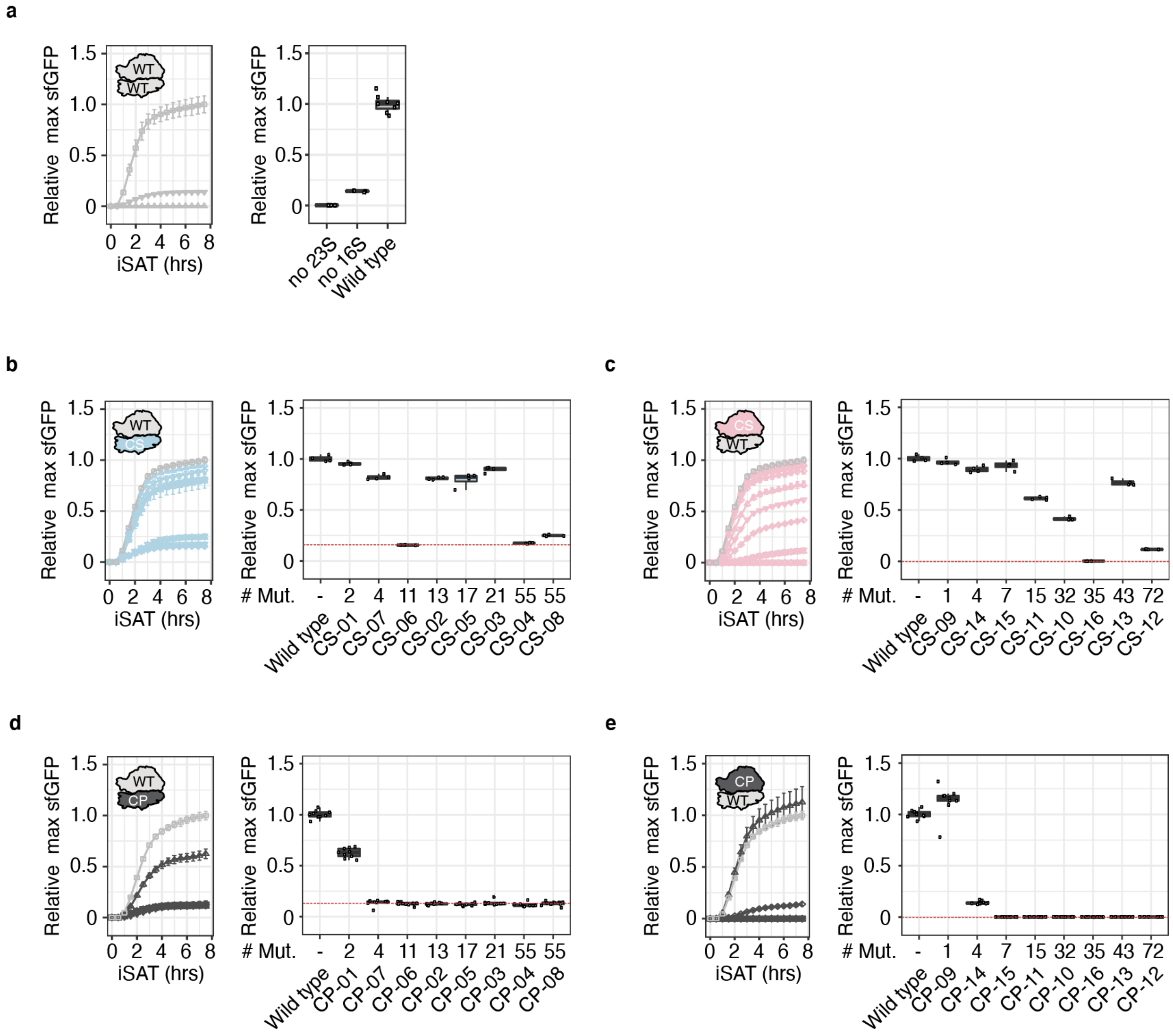


**Supplementary Figure 1: sfGFP expression of 16S and 23S rRNA designs during iSAT.** (**a**). Community scientist-designed (CS) 16S rRNA (**b**) and 23S rRNA (**c**) designs. Computationally predicted (CP) 16S rRNA (**d**) and 23S rRNA (**e**). sfGFP expression in iSAT was determined by fluorescence over the course of 8 hours and normalized to the maximum sfGFP made by the wild type ribosome. Time course data are shown as mean ± s.d. on the left of each panel and the relative max sfGFP generated by each design as boxplots on its right side. Error bars represent s.d.; n ≥ 4. Dotted red line indicates background activity arising from the extract. Mut: mutations; WT: wild type.

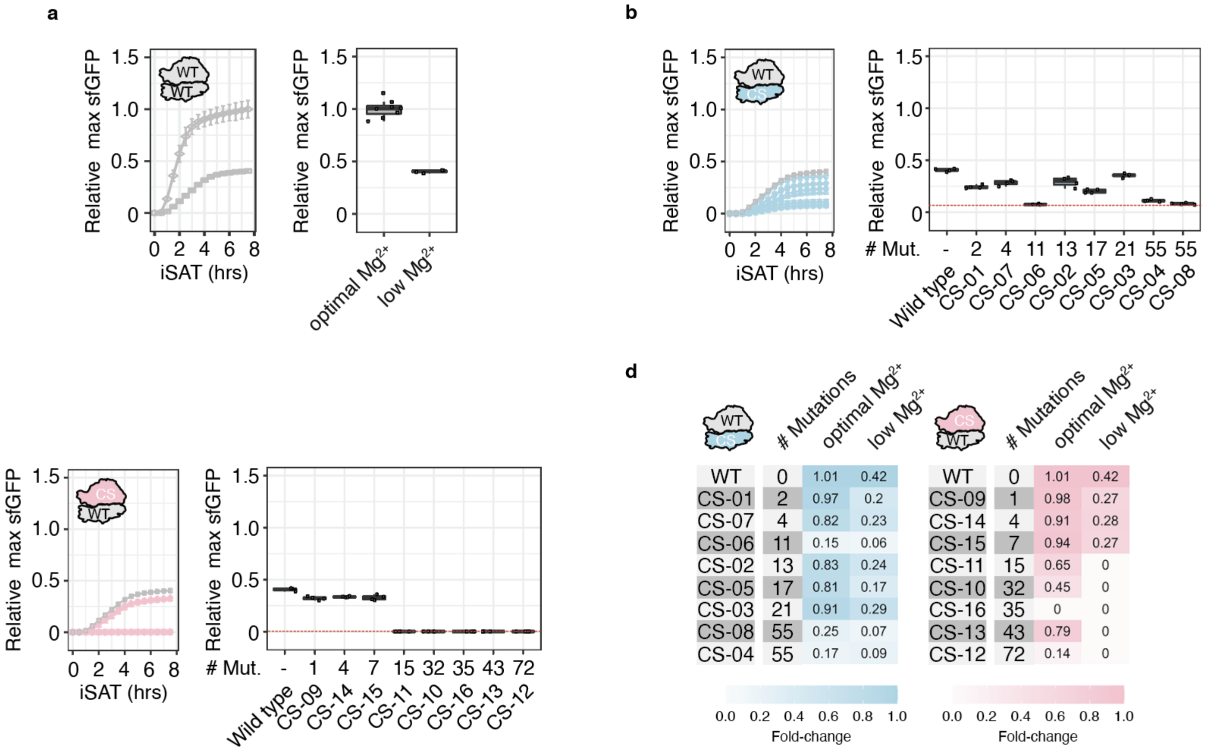


**Supplementary Figure 2: Eterna rRNAs can enable cell-free translation in folding stress conditions.** (**a**) Relative sfGFP expression of wild type ribosomes under optimal and low (3.75 mM)-magnesium ($Mg^{2+}$) iSAT conditions. Performance of Eterna 16S rRNAs (**b**) and 23S rRNAs (**c**) at low magnesium iSAT conditions. (**d**) Heatmap illustrating number of rRNA mutations and relative maximum sfGFP expression in optimal (7.5 mM) and low (3.75 mM)-magnesium iSAT reactions of pT7-rrnB-16S (left) and pT7-rrnB-23S (right) wild type and variants designed by community scientists (CS). sfGFP expression was determined by fluorescence over 8 hours and normalized to the maximum sfGFP of pT7-rrnB-WT at optimal iSAT conditions. Max sfGFP made by each design is shown as means normalized to pT7-rrnB-wild type activity at optimal iSAT conditions. Error bars represent s.d.; n ≥ 4. Dotted red line indicates background activity arising from the extract. Mut: mutations; WT: wild type.

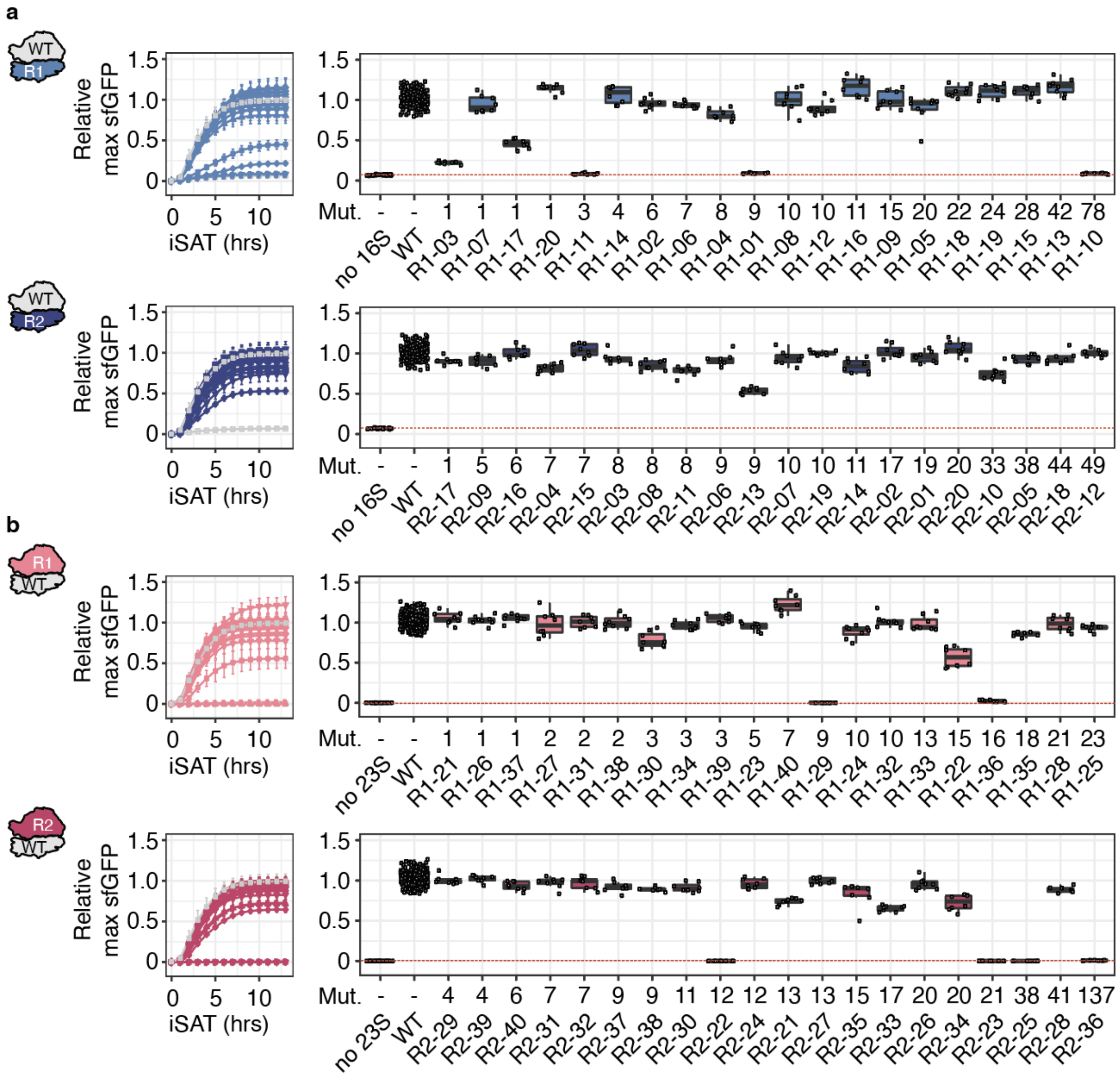


**Supplementary Figure 3: iSAT time courses of Round (R1) and Round 2 (R2) Eterna designed ribosomes.** sfGFP expression of pT7-rrnB-R1 and pT7-rrnB-R2 16S rRNA (**a**) and pT7-rrnB-R1 and pT7-rrnB-R2 23S rRNA (**b**) designs for 16-hour iSAT reactions. sfGFP expression in iSAT was determined by fluorescence and normalized to max sfGFP of pT7-rrnB-wild type. Error bars represent s.d.; n ≥ 3. Dotted red line indicates background activity arising from the extract. Mut: mutations, R1: round 1, R2: round 2, WT: wild type.

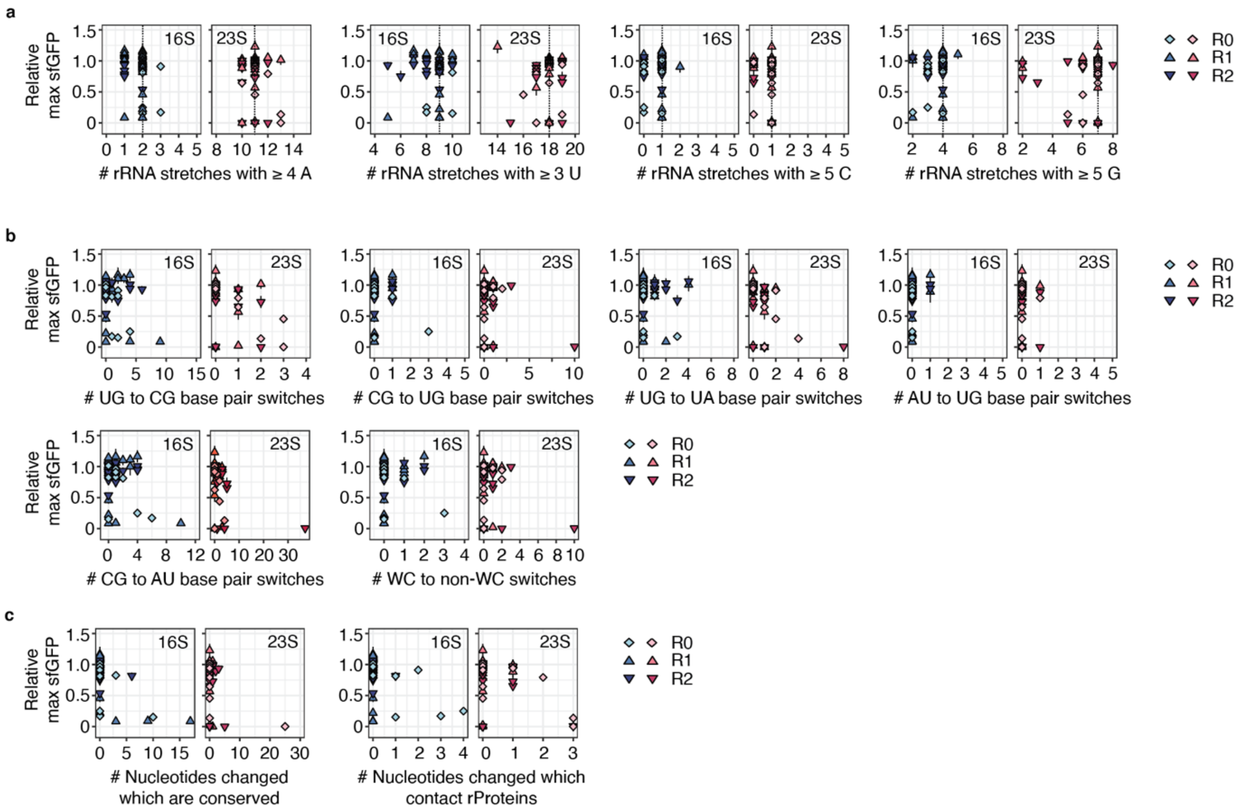


**Supplementary Figure 4: Community scientists followed and combined different strategies to improve rRNA performance.** Relative maximum sfGFP expression made in iSAT reactions by each design was plotted against the instances of (**a**) stretches of consecutive identical nucleotides, (**b**) altered base pairing in rRNA secondary structures, or (**c**) changes in conserved nucleotides or nucleotides which contact rProteins. Data are shown from the "pilot round" (R0) and round 1 (R1) and round 2 (R2) as mean ± s.d.; n ≥ 3. Dotted line in (a) indicates wild type value. rProteins: ribosomal proteins; WC: Watson-Crick.

WT / R1

| | # Mutations | Low Mg$^{2+}$ / 37°C | Low Mg$^{2+}$ / 30°C |
|---|---|---|---|
| WT | 0 | 0.41 | 0.33 |
| R1-02 | 6 | 0.39 | 0.3 |
| R1-06 | 7 | 0.43 | 0.32 |
| R1-04 | 8 | 0.26 | 0.19 |
| R1-12 | 10 | 0.36 | 0.27 |
| R1-08 | 10 | 0.44 | 0.31 |
| R1-16 | 11 | 0.41 | 0.33 |
| R1-09 | 15 | 0.41 | 0.3 |
| R1-05 | 20 | 0.38 | 0.3 |
| R1-18 | 22 | 0.34 | 0.32 |
| R1-19 | 24 | 0.26 | 0.28 |
| R1-15 | 28 | 0.41 | 0.31 |
| R1-13 | 42 | 0.42 | 0.35 |

WT / R2

| | # Mutations | Low Mg$^{2+}$ / 37°C | Low Mg$^{2+}$ / 30°C |
|---|---|---|---|
| WT | 0 | 0.41 | 0.33 |
| R2-16 | 6 | 0.42 | 0.37 |
| R2-15 | 7 | 0.47 | 0.37 |
| R2-03 | 8 | 0.4 | 0.11 |
| R2-06 | 9 | 0.41 | 0.11 |
| R2-07 | 10 | 0.36 | 0.09 |
| R2-19 | 10 | 0.42 | 0.35 |
| R2-02 | 17 | 0.41 | 0.32 |
| R2-01 | 19 | 0.44 | 0.11 |
| R2-20 | 20 | 0.38 | 0.35 |
| R2-05 | 38 | 0.37 | 0.13 |
| R2-18 | 44 | 0.4 | 0.26 |
| R2-12 | 49 | 0.39 | 0.37 |

R1 / WT

| | # Mutations | Low Mg$^{2+}$ / 37°C | Low Mg$^{2+}$ / 30°C |
|---|---|---|---|
| WT | 0 | 0.41 | 0.33 |
| R1-40 | 7 | 0.37 | 0.31 |
| R1-32 | 10 | 0.02 | 0.03 |
| R1-24 | 10 | 0 | 0 |
| R1-33 | 13 | 0.21 | 0.19 |
| R1-35 | 18 | 0.07 | 0.03 |
| R1-28 | 21 | 0.02 | 0.02 |
| R1-25 | 23 | 0.16 | 0.18 |

R2 / WT

| | # Mutations | Low Mg$^{2+}$ / 37°C | Low Mg$^{2+}$ / 30°C |
|---|---|---|---|
| WT | 0 | 0.41 | 0.33 |
| R2-40 | 6 | 0.15 | 0.12 |
| R2-32 | 7 | 0.22 | 0.32 |
| R2-31 | 7 | 0.05 | 0.36 |
| R2-37 | 9 | 0.09 | 0.22 |
| R2-38 | 9 | 0.3 | 0.17 |
| R2-30 | 11 | 0.22 | 0.17 |
| R2-24 | 12 | 0.32 | 0.28 |
| R2-27 | 13 | 0.32 | 0.18 |
| R2-26 | 20 | 0.28 | 0.38 |
| R2-28 | 41 | 0.43 | 0.1 |

**Supplementary Figure 5: Eterna designs are robust across diverse folding stress conditions *in vitro***. Data are replotted from Main Figure 4a, but with numerical values listed. sfGFP expression in iSAT was determined by fluorescence and normalized to max sfGFP of pT7-rrnB-wild type at optimal iSAT conditions. Data are shown as mean; n ≥ 3. R1: round 1, R2: round 2, WT: wild type.

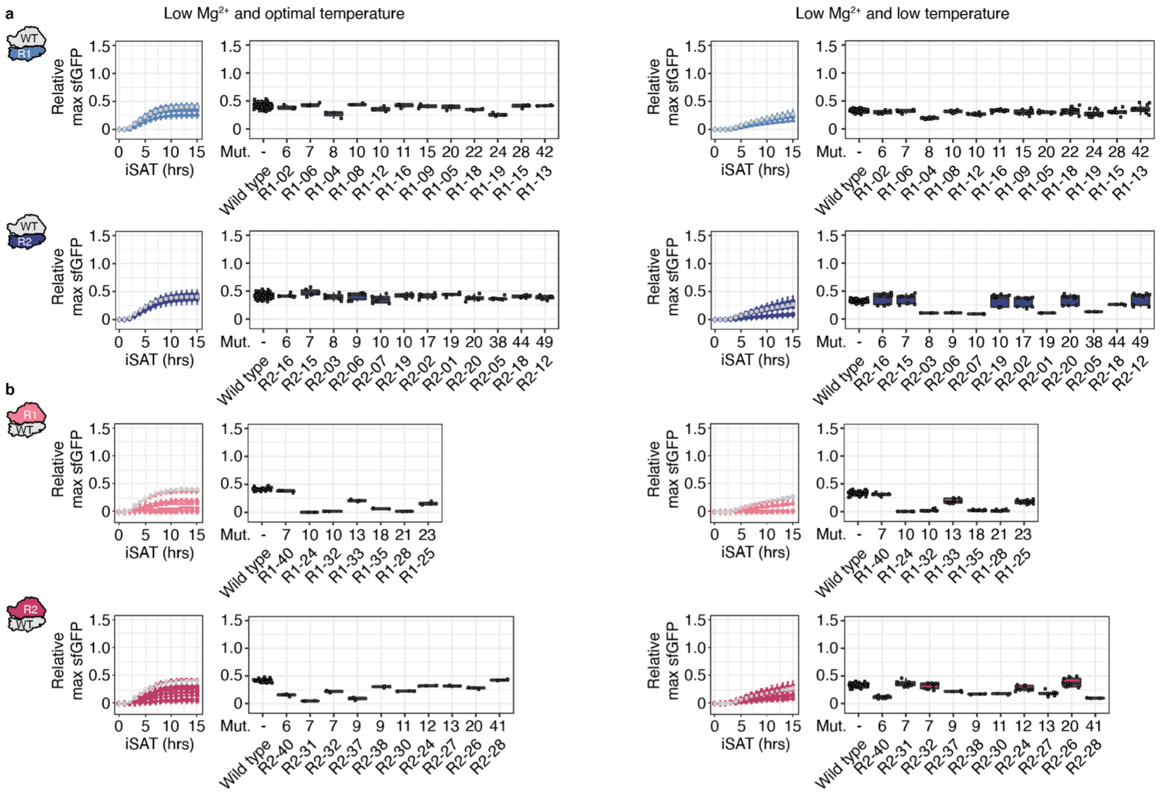

**Supplementary Figure 6: iSAT kinetics and maximum yields for Eterna designed ribosomes under folding stress.** sfGFP expression of (**a**) pT7-rrnB-16S R1 and R2 designs and (**b**) pT7-rrnB-23S R1 and R2 designs in iSAT at low $Mg^{2+}$ concentration (3.75 mM) and optimal temperature (37º C) (left panels) and low $Mg^{2+}$ concentration (3.75 mM) and low temperature (30º C) (right panels). sfGFP expression was determined in 15 hour iSAT reactions by fluorescence and normalized to max sfGFP of pT7-rrnB-wild type at optimal iSAT conditions. Data are shown as mean. Error bars represent s.d.; n ≥ 3. These data were used to generate the heat maps in Main Figure 3 and Suppl. Figure 5. Mut: mutations; R1: round 1, R2: round 2, WT: wild type.

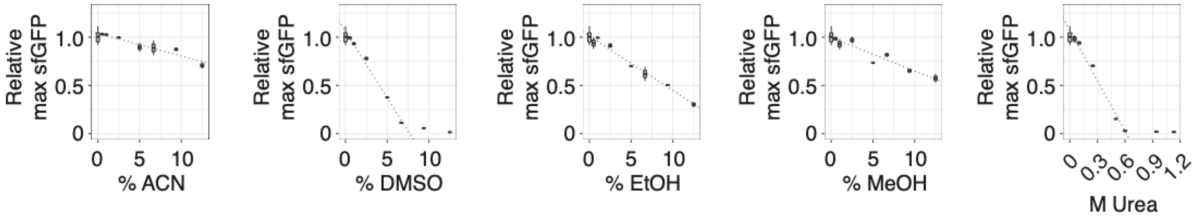


**Supplementary Figure 7: Influence of solvents on iSAT reactions.** sfGFP expression of pT7-rrnB-WT in iSAT in the presence of increasing concentrations of solvents. Solvents are shown based on volume percent. ACN: acetonitrile, DMSO: dimethylsulfoxide, EtOH: ethanol, MeOH: methanol. Data are shown as boxplots. Error bars represent s.d.; n ≥ 3.

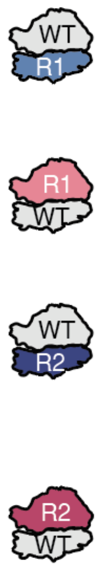


| | 9.7% ACN | 12.5% ACN | 3.3% DMSO | 6.7% DMSO | 3.3% EtOH | 6.7% EtOH | 6.7% MeOH | 9.7% MeOH | 12.5% MeOH | 0.33 M Urea | pH 6.4 | pH 7.2 | pH 8.0 | pH 8.8 |
|---|---|---|---|---|---|---|---|---|---|---|---|---|---|---|
| | Solvents | | | | | | | | | | pH | | | |
| WT | 0.82 | 0.76 | 0.61 | 0.14 | 0.83 | 0.65 | 0.81 | 0.68 | 0.58 | 0.5 | 0.58 | 0.85 | 0.35 | 0.24 |
| R1-05 | 0.58 | 0.57 | 0.44 | 0.23 | 0.56 | 0.53 | 0.82 | 0.64 | 0.72 | 0.43 | 0.41 | 0.59 | 0.1 | 0.04 |
| R1-09 | 0.79 | 0.86 | 0.62 | 0.28 | 0.88 | 0.62 | 0.75 | 0.74 | 0.6 | 0.5 | 0.64 | 0.85 | 0.18 | 0.12 |
| R1-13 | 0.72 | 0.72 | 0.5 | 0.24 | 0.84 | 0.82 | 0.78 | 0.91 | 0.93 | 0.5 | 0.43 | 0.69 | 0.14 | 0.09 |
| R1-15 | 0.75 | 0.81 | 0.58 | 0.24 | 0.94 | 0.71 | 0.79 | 0.78 | 0.85 | 0.48 | 0.53 | 0.75 | 0.18 | 0.11 |
| R1-18 | 0.58 | 0.54 | 0.46 | 0.1 | 0.66 | 0.48 | 0.7 | 0.55 | 0.56 | 0.42 | 0.31 | 0.45 | 0.04 | 0.01 |
| R1-25 | 0.7 | 1.06 | 0.56 | 0.09 | 0.93 | 1.04 | 0.91 | 0.69 | 0.36 | 0.32 | 0.47 | 0.6 | 0.07 | 0.04 |
| R1-33 | 1.08 | 0.91 | 0.72 | 0.2 | 1 | 1.2 | 0.99 | 0.87 | 0.4 | 0.45 | 0.53 | 0.65 | 0.1 | 0.07 |
| R1-40 | 0.76 | 0.68 | 0.42 | 0.12 | 0.6 | 0.66 | 0.88 | 0.38 | 0.21 | 0.51 | 0.39 | 0.5 | 0.07 | 0.05 |
| R2-02 | 1.06 | 0.89 | 0.64 | 0.16 | 1.03 | 1.07 | 0.7 | 0.67 | 0.35 | 0.52 | 0.52 | 0.65 | 0.21 | 0.15 |
| R2-05 | 1.05 | 0.83 | 0.58 | 0.17 | 0.97 | 1.16 | 0.94 | 0.73 | 0.35 | 0.48 | 0.44 | 0.58 | 0.14 | 0.09 |
| R2-12 | 0.97 | 0.75 | 0.41 | 0.11 | 0.63 | 1.09 | 0.85 | 0.71 | 0.56 | 0.49 | 0.37 | 0.51 | 0.07 | 0.05 |
| R2-18 | 0.12 | 0.18 | 0.09 | 0.04 | 0.04 | 0.16 | 0.81 | 0.05 | 0.17 | 0.49 | 0.5 | 0.68 | 0.16 | 0.08 |
| R2-20 | 0.42 | 0.36 | 0.18 | 0.05 | 0.19 | 0.45 | 0.74 | 0.26 | 0.25 | 0.47 | 0.21 | 0.28 | 0.02 | 0.01 |
| R2-24 | 0.29 | 0.46 | 0.17 | 0.05 | 0.13 | 0.25 | 0.74 | 0.18 | 0.14 | 0.35 | 0.22 | 0.3 | 0.01 | 0.01 |
| R2-26 | 0.77 | 0.85 | 0.55 | 0.13 | 0.71 | 0.67 | 0.8 | 0.57 | 0.38 | 0.51 | 0.48 | 0.67 | 0.12 | 0.12 |
| R2-27 | 0.48 | 0.56 | 0.25 | 0.09 | 0.29 | 0.34 | 0.73 | 0.28 | 0.2 | 0.38 | 0.32 | 0.46 | 0.01 | 0.01 |
| R2-28 | 0.76 | 1.03 | 0.52 | 0.12 | 0.76 | 0.69 | 0.74 | 0.45 | 0.2 | 0.44 | 0.48 | 0.62 | 0.09 | 0.07 |
| R2-38 | 0.77 | 0.71 | 0.55 | 0.15 | 0.88 | 0.66 | 0.65 | 0.46 | 0.25 | 0.44 | 0.39 | 0.76 | 0.16 | 0.16 |

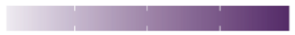


**Supplementary Figure 8: Eterna designs are robust across diverse *in vitro* stress conditions**. Data are replotted from Main Figure 4b, but with numerical values listed. sfGFP expression in iSAT was determined by fluorescence and normalized to the maximum sfGFP of pT7- rrnB-WT at optimal iSAT conditions. Data are shown as mean; n ≥ 3. ACN: acetonitrile, DMSO: dimethylsulfoxide, EtOH: ethanol, MeOH: methanol, HEPES: (4-(2-hydroxyethyl)-1-piperazineethanesulfonic acid), MES: 2-(N-morpholino)ethanesulfonic acid, Tris: tris(hydroxymethyl)aminomethane, Mut: mutations, R1: round 1, R2: round 2, WT: wild type. 16S rRNA and 23S rRNA designs whose names have been highlighted represent the most diverse and robust 16S rRNA and 23S rRNA designs per round.

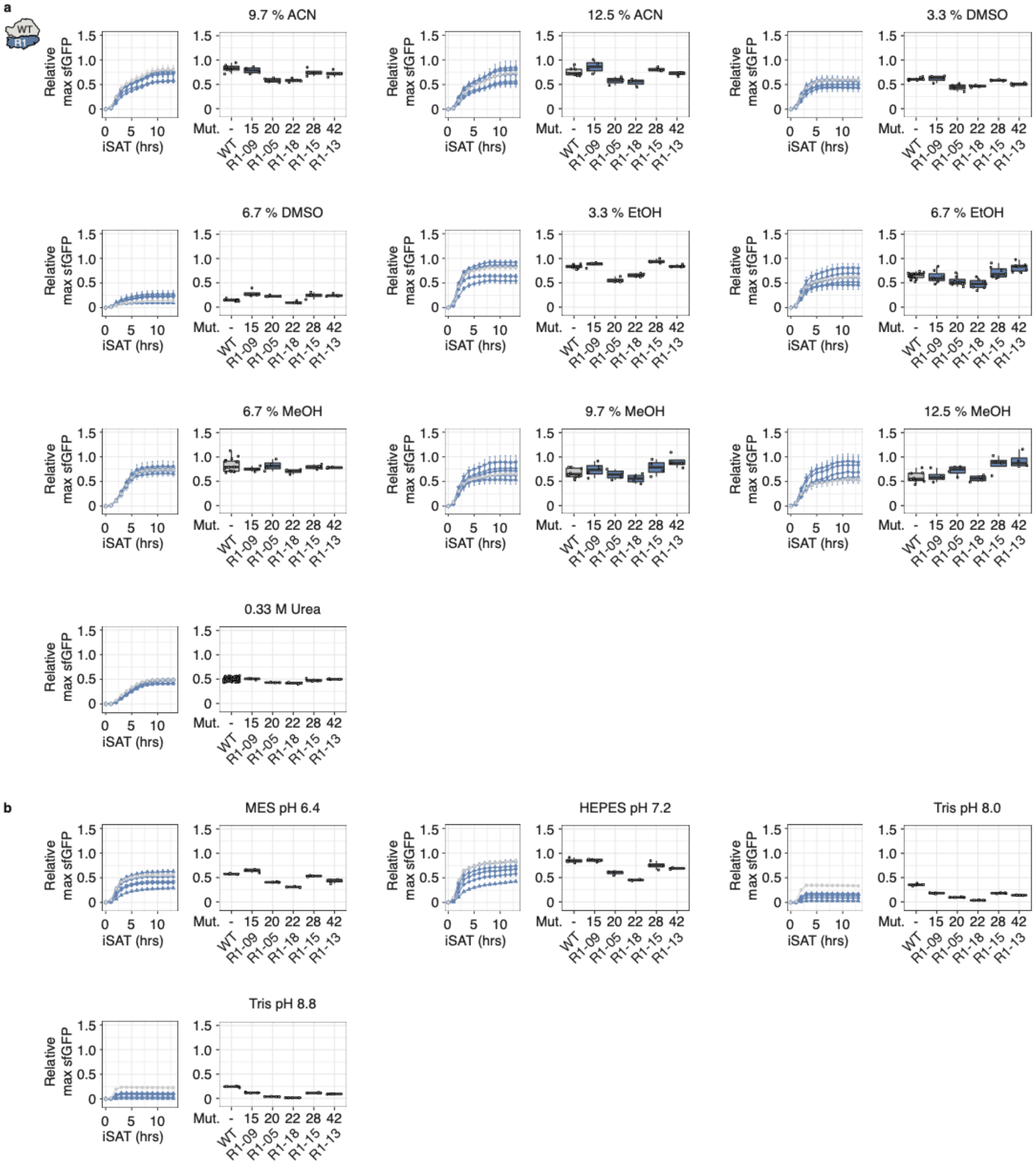


**Supplementary Figure 9: Influence of solvents and pH on iSAT reactions of selected R1 16S rRNA designs.** (**a**) Solvents (v/v %). (**b**) pH. Time course data are shown as mean ± s.d. Maximum sfGFP expression was determined in iSAT reactions by fluorescence and normalized to max sfGFP of pT7-rrnB-wild type at optimal iSAT conditions. Maximal sfGFP expression data are presented as boxplots. Error bars represent s.d.; n ≥ 3. ACN: acetonitrile, DMSO: dimethylsulfoxide, EtOH: ethanol, MeOH: methanol, HEPES: (4-(2-hydroxyethyl)-1-piperazineethanesulfonic acid), MES: 2-(N-morpholino)ethanesulfonic acid, Tris: tris(hydroxymethyl)aminomethane, Mut: mutations, R1: round 1, WT: wild type.

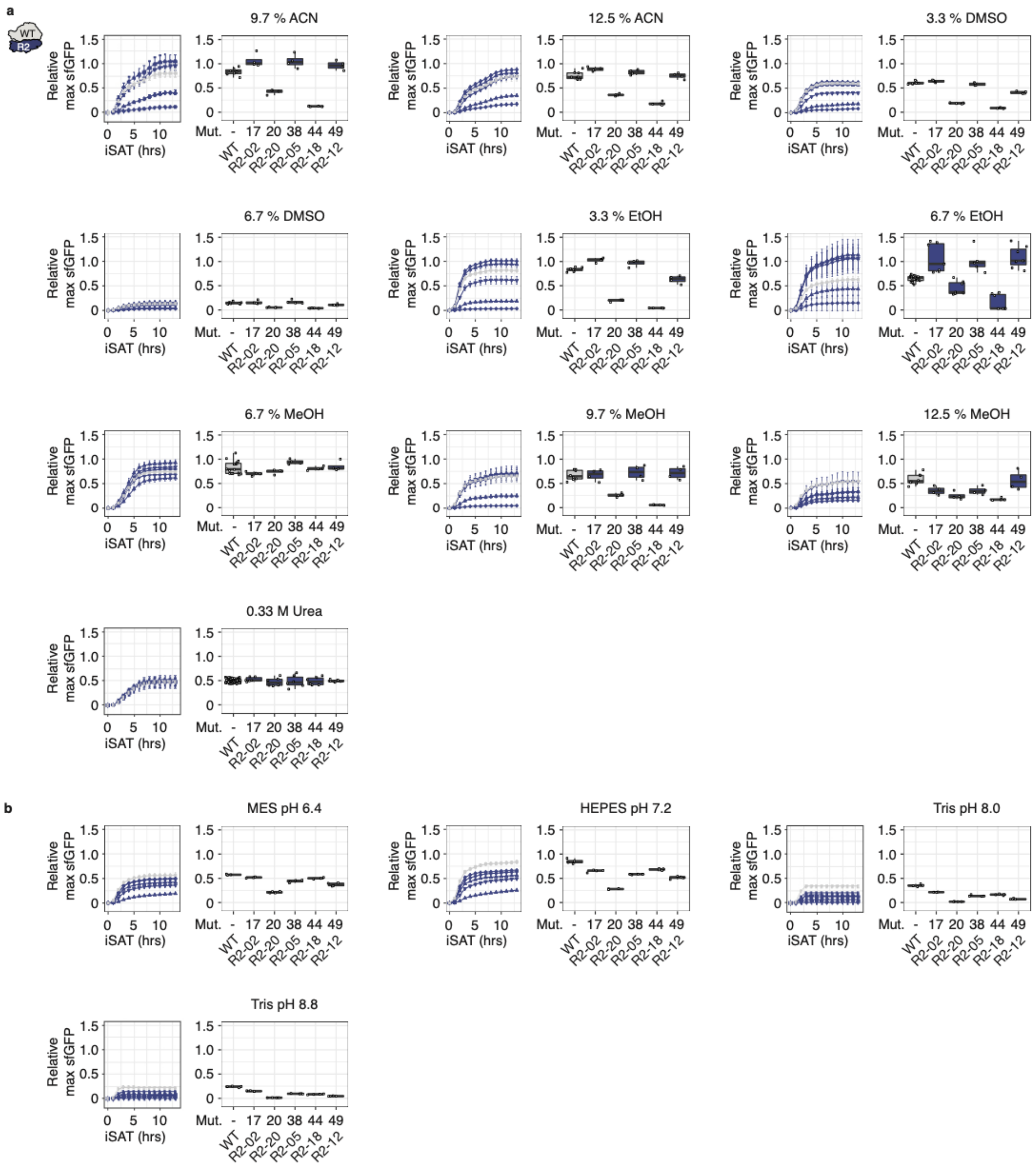


**Supplementary Figure 10: Influence of solvents and pH on iSAT reactions of selected R2 16S rRNA designs.** (**a**) Solvents (v/v %). (**b**) pH. Time course data are shown as mean ± s.d. Maximum sfGFP expression was determined in iSAT reactions by fluorescence and normalized to max sfGFP of pT7-rrnB-wild type at optimal iSAT conditions. Maximal sfGFP expression data are presented as boxplots. Error bars represent s.d.; n ≥ 3. ACN: acetonitrile, DMSO: dimethylsulfoxide, EtOH: ethanol, MeOH: methanol, HEPES: (4-(2-hydroxyethyl)-1-piperazineethanesulfonic acid), MES: 2-(N-morpholino)ethanesulfonic acid, Tris: tris(hydroxymethyl)aminomethane, Mut: mutations, R2: round 1, WT: wild type

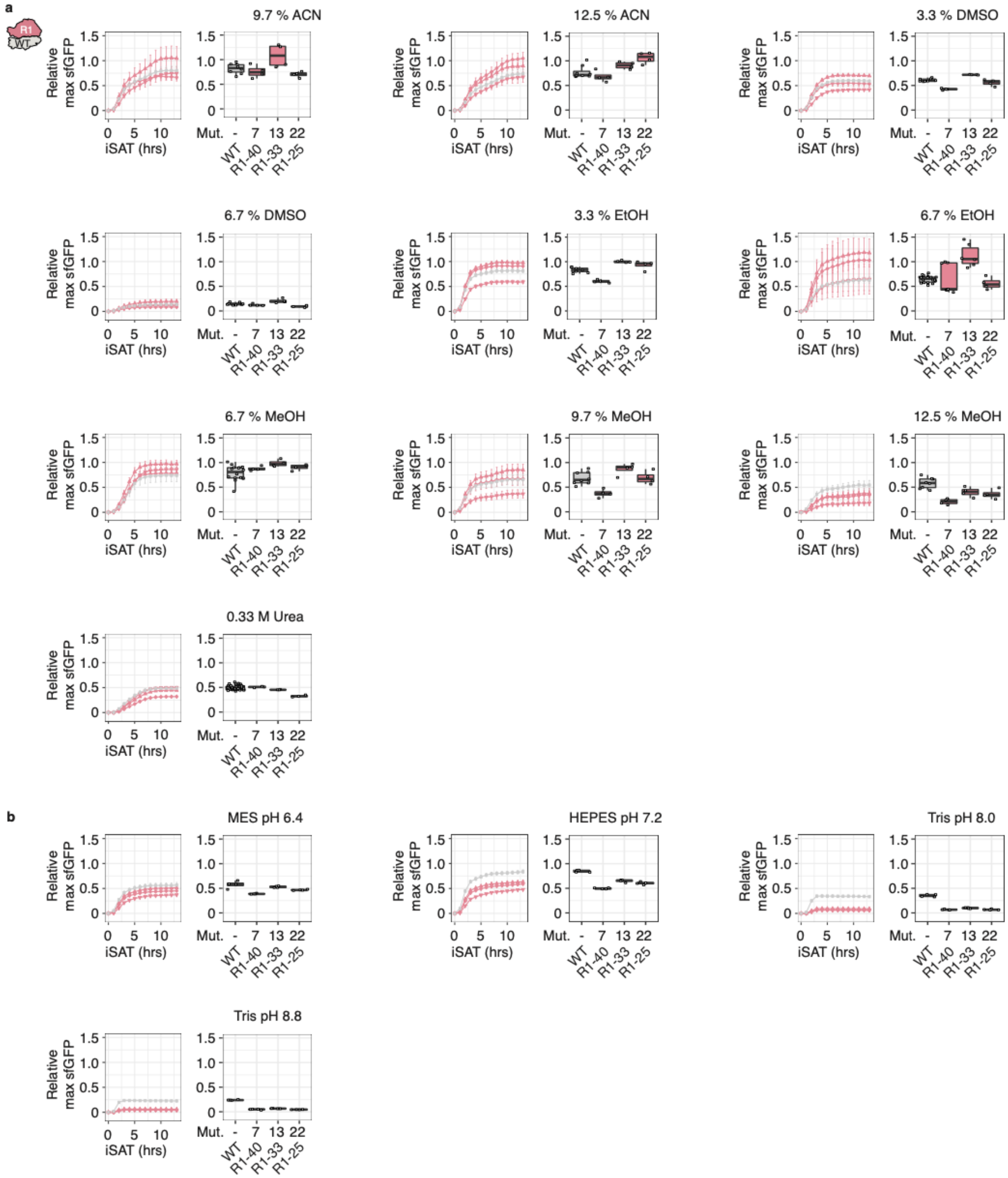


**Supplementary Figure 11: Influence of solvents and pH on iSAT reactions of selected R1 23S rRNA designs.** (**a**) Solvents (v/v %). (**b**) pH. Time course data are shown as mean ± s.d. Maximum sfGFP expression was determined in iSAT reactions by fluorescence and normalized to max sfGFP of pT7-rrnB-wild type at optimal iSAT conditions. Maximal sfGFP expression data are presented as boxplots. Error bars represent s.d.; n ≥ 3. ACN: acetonitrile, DMSO: dimethylsulfoxide, EtOH: ethanol, MeOH: methanol, HEPES: (4-(2-hydroxyethyl)-1-piperazineethanesulfonic acid), MES: 2-(N-morpholino)ethanesulfonic acid, Tris: tris(hydroxymethyl)aminomethane, Mut: mutations, R1: round 1, WT: wild type.

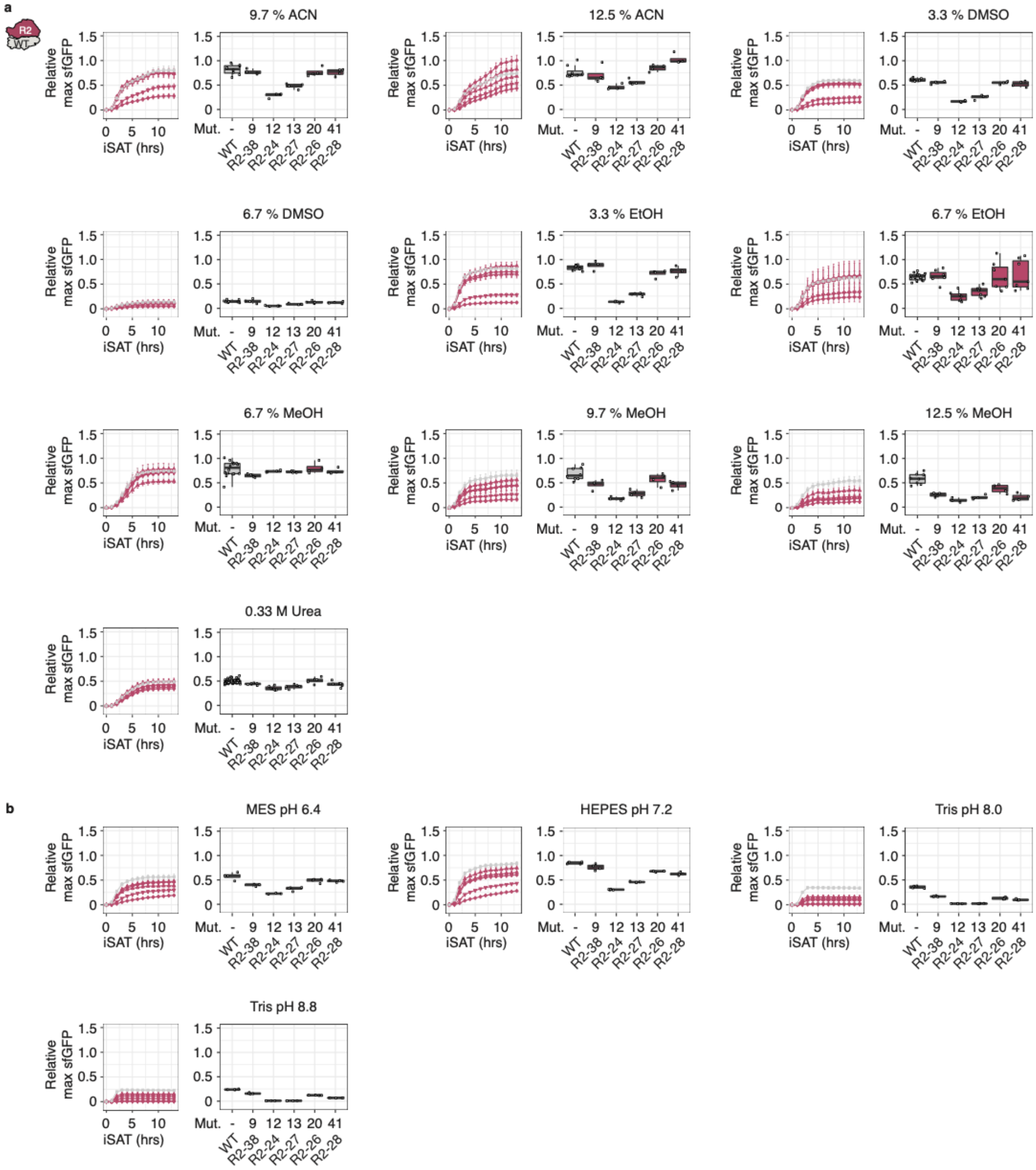


**Supplementary Figure 12: Influence of solvents and pH on iSAT reactions of selected R2 23S designs.** (**a**) Solvents (v/v %). (**b**) pH. Time course data are shown as mean ± sd. Maximum sfGFP expression was determined in iSAT reactions by fluorescence and normalized to max sfGFP of pT7-rrnB-wild type at optimal iSAT conditions. Maximal sfGFP expression data are presented as boxplots. Error bars represent s.d.; n ≥ 3. ACN: acetonitrile, DMSO: dimethylsulfoxide, EtOH: ethanol, MeOH: methanol, HEPES: (4-(2-hydroxyethyl)-1-piperazineethanesulfonic acid), MES: 2-(N-morpholino)ethanesulfonic acid, Tris: tris(hydroxymethyl)aminomethane, Mut: mutations, R2: round 2, WT: wild type.

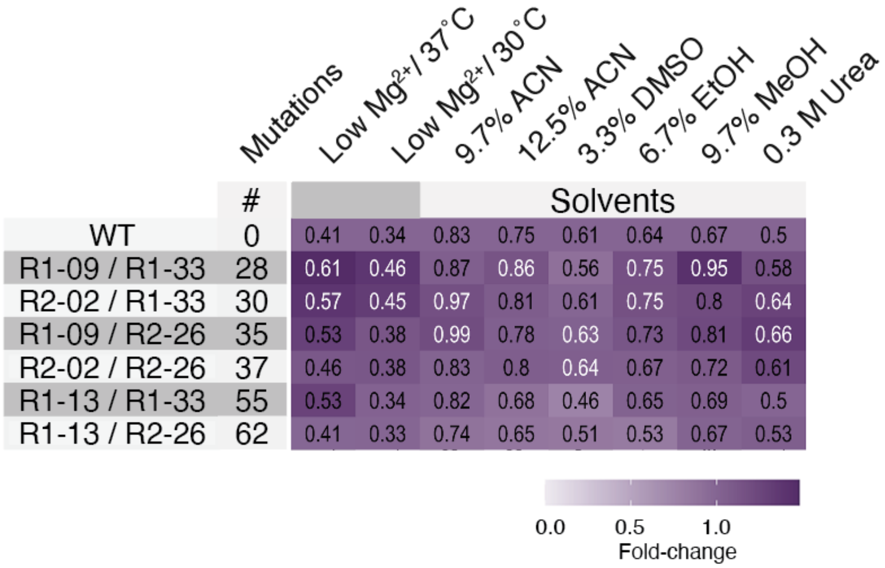


**Supplementary Figure 13: Eterna designs are robust across diverse *in vitro* stress conditions** (i.e., solvents). Data are replotted from Main Figure 5a, but with numerical values listed. sfGFP expression in iSAT was determined by fluorescence and normalized to maximum sfGFP of pT7- rrnB-wild type at optimal iSAT conditions. Data are shown as mean; n ≥ 3. ACN: acetonitrile, DMSO: dimethylsulfoxide, EtOH: ethanol, MeOH: methanol, WT: wild type.

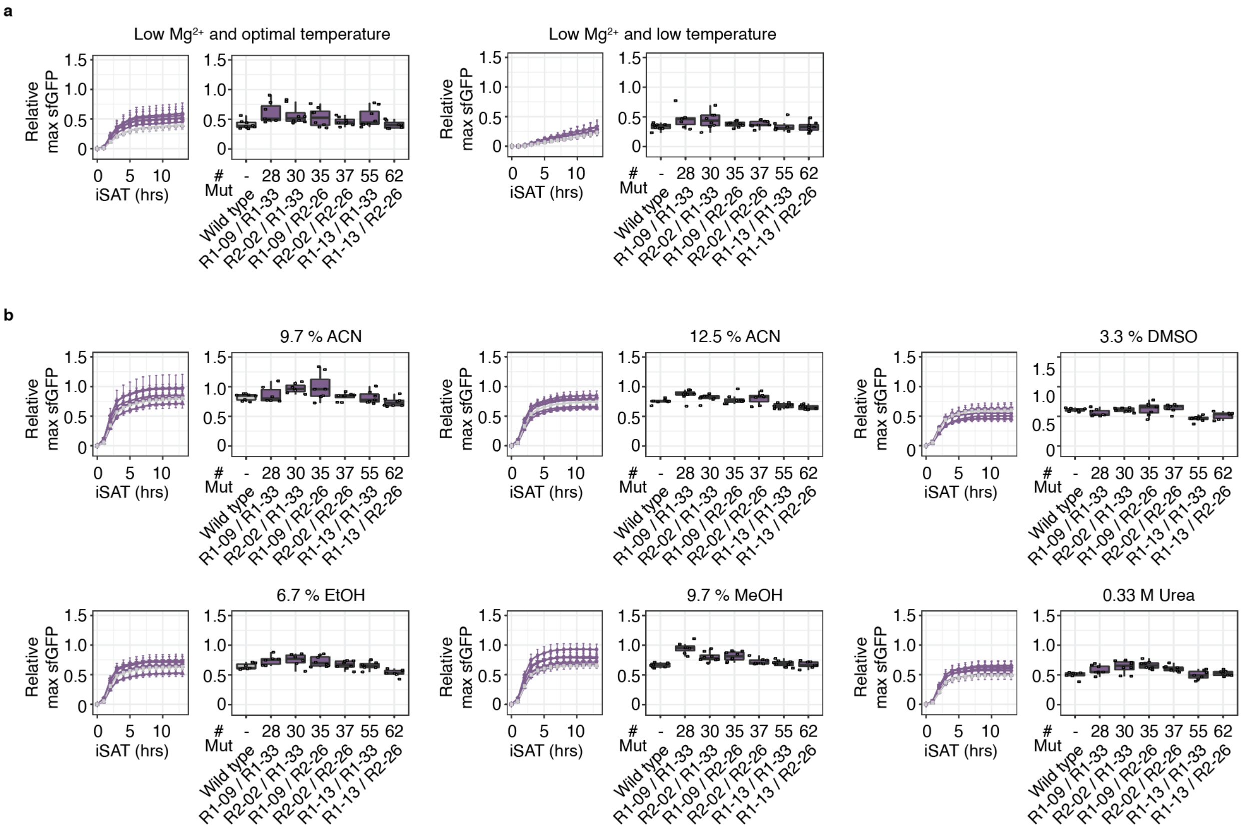


**Supplementary Figure 14: Influence of *in vitro* solvent conditions on iSAT reactions of Eterna ribosomes.** (**a**) Relative sfGFP expression of ribosomes with both 16S rRNA and 23S rRNA designs under iSAT conditions at low (3.75 mM)-magnesium ($Mg^{2+}$) and optimal or low (3.75 mM)-magnesium ($Mg^{2+}$) and low temperature. (**b**) Solvents (v/v %). Time course data are shown as mean ± s.d. Maximum sfGFP expression was determined in iSAT reactions by fluorescence and normalized to max sfGFP of pT7-rrnB-wild type at optimal iSAT conditions. Maximal sfGFP expression data are presented as boxplots. Error bars represent s.d.; n ≥ 3. ACN: acetonitrile, DMSO: dimethylsulfoxide, EtOH: ethanol, MeOH: methanol, Mut: mutations.

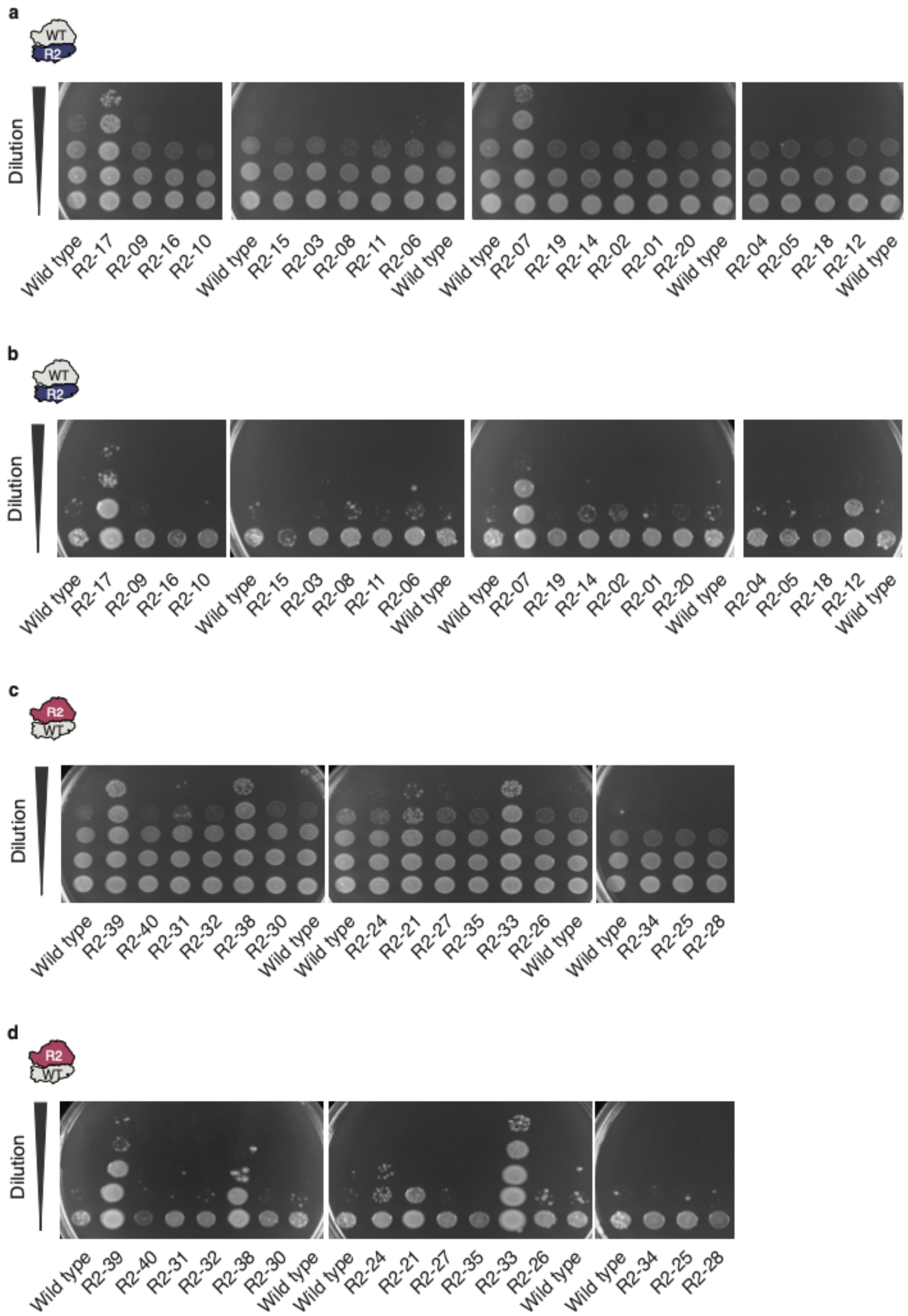


**Supplementary Figure 15: R2 Eterna designs support life.** (**a-d**) Un-cut images of spotted SQ171fg cells growing with pL-rrnB-wild type and pL-rrnB-R2 16S rRNA (**a, b**) and 23S rRNA (**c, d**) designs imaged after 24 hours at 37 °C (**a, c**) or 72 hours at 30 °C (**b, d**). Stationary cells were diluted to an OD600 = 1, diluted stepwise 1:10, and spotted onto LB + Carb100 plates. Data are representative of n ≥ 3. R2: round 2, WT: wild type.

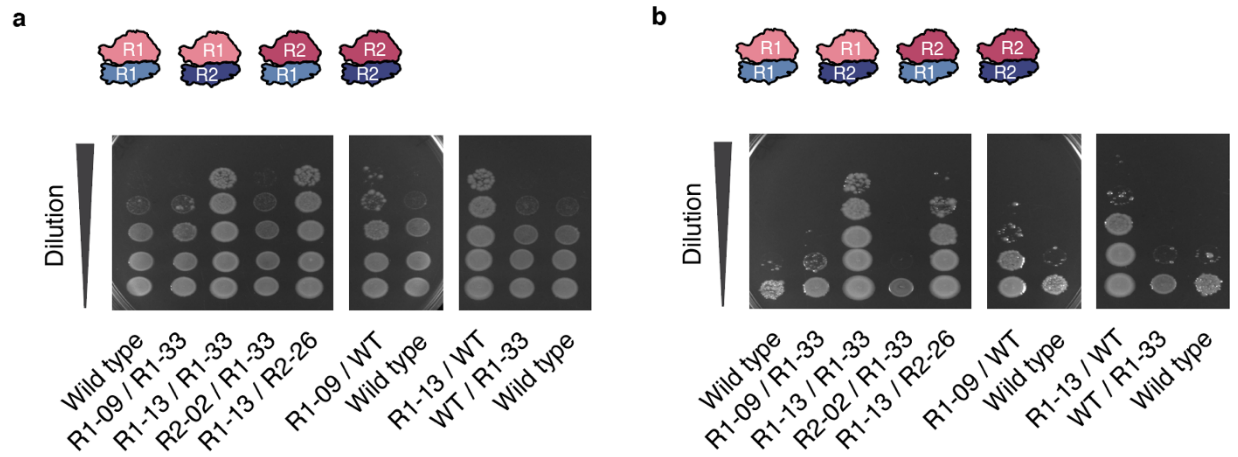


**Supplementary Figure 16: Combinatorial Eterna designs support life.** (**a-b**) Un-cut images of spotted SQ171fg cells growing with pL-rrnB-wild type and pL-rrnB-Combinations imaged after 24 hours at 37 °C (**a**) or 72 hours at 30 °C (**b**). Stationary cells were diluted to an OD600 = 1, diluted stepwise 1:10, and spotted onto LB + Carb100 plates. Data are representative of n ≥ 3. R1: round 1, R2: round 2, WT: wild type.
